# Functional human neurospheroids recapitulate key features of cortical complexity

**DOI:** 10.64898/2026.03.09.710475

**Authors:** Giulia Parodi, Giorgia Zanini, Linda Collo, Donatella Di Lisa, Cecilia Beccari, Michela Chiappalone, Sergio Martinoia

## Abstract

**Background:** Three-dimensional *in vitro* neuronal cultures have emerged as promising platforms for modelling human brain function and disease under controlled conditions. However, their ability to recapitulate *in vivo*-like complexity and rich dynamics remains underexplored. In this study, we developed and characterized neurospheroids derived from human induced pluripotent stem cells (hiPSCs) to investigate how key features — three-dimensionality, cellular heterogeneity, and modular organization — contribute to replicating brain-like network dynamics.

**Methods:** We engineered neurospheroids with varying excitatory/inhibitory ratios and assembled them into modular constructs (assembloids), evaluating their electrophysiological activity using high-density micro-electrode arrays. We assessed spontaneous and evoked activity through established metrics of dynamical richness and perturbational complexity.

**Results:** Our findings show that three-dimensionality and modularity significantly enhance the richness and complexity of network activity, approaching levels observed *in vivo*, while cellular heterogeneity contribute to functional aspects of the network, such as nuanced activity patterns, dynamical variability, and modular coordination.

**Conclusions:** Our work highlights the critical role of spatial organization in reproducing brain-like activity and provide a foundation for future studies using patient-derived neurospheroids to model disease-specific dynamics.

## Introduction

*In vitro* models offer a simplified yet powerful approach for studying the human brain, enabling the investigation of neuronal diseases under controlled conditions. Capturing key *in vivo* features such as three-dimensional (3D) structure, cellular heterogeneity, and modular organization is crucial for developing reliable experimental *in vitro* models that support consistent and reproducible research.

Nowadays, 3D *in vitro* neuronal cultures allow to replicate the intricate spatial structure of the brain, providing better insights of the neuronal development, function, and disease mechanisms^1,2^. Among several possibilities, brain organoids and neurospheroids are widely leveraged as 3D neuronal cultures^3^. Indeed, the self-organization of the neurons in the 3D structure can recapitulate key features of the human brain^4^, mimicking its developmental processes^5,6^. For these reasons, 3D models constitute a valuable platform for exploring neuronal growth and interaction in both physiological and pathological conditions, e.g., neurodevelopmental disorders^7,8^, as well as for drug screening^9^ or for exploring personalized medicine approaches^10^. Furthermore, they represent a useful environment for studying basic neuronal biology and synaptic functions^11^.

Since their first introduction in 1962^12^, 3D *in vitro* cultures have rapidly gained widespread use, especially in the last years^13,14^. More recently, substantial progress has been accomplished through the use of human-derived cells^15–17^, which more accurately replicates the *in vivo* condition^18^. This approach enables a deeper understanding of human disease mechanisms and progression, supports drug discovery and testing, and facilitates the development of cell replacement therapy^4^. In parallel with the development of 3D experimental models, 2D models evolved to incorporate new attributes such as heterogeneity and modularity, both in rodent^19–21^ and in human induced pluripotent stem cells (hiPSCs)-derived neuronal cultures^22–26^. Despite the evolution and the current availability of the above-mentioned *in vitro* systems, the properties of hiPSC-derived 3D models, in particular neurospheroids, have received limited attention in the scientific literature. Crucially, investigations into heterogeneous neurospheroids with a well-defined excitation/inhibition (E/I) balance, i.e., a key aspect of neural network function, remain largely unexplored, leaving a significant gap in our understanding of their full potential.

In this work, we aim to fill this scientific gap, necessary to answer important key questions in the field, which have so far remained unexplored: to what extent are human-derived neurospheroids able to recapitulate the intrinsic rich dynamics and complexity of the *in vivo* cortex? Are these models ‘better’ than classical 2D cultures? Are the features of three-dimensionality, heterogeneity, and modularity all necessary to reproduce cortical brain complexity?

To respond to these questions, we first designed a simple, reliable and repeatable technique to create 3D experimental models based on neuronal networks derived from hiPSCs, including distinctive features such as heterogeneity and modularity. Specifically, by capitalizing on our experience on 2D human-derived culture^27,28^, we realized engineered neurospheroids composed of different excitation/inhibition (E/I) ratios: we created homogeneous excitatory or inhibitory neurospheroids in which only glutamatergic or GABAergic neurons were included, respectively, while the heterogeneous configuration consisted in 75% of glutamatergic neurons and 25% of GABAergic ones^29^. To implement modularity, we realized assembloids^30–32^, i.e., two neurospheroids brought into contact. We then evaluated the appropriate dimension of the neurospheroids to ensure good adhesion to the recording substrate (2304 or 4096 micro-electrodes of a high-density Micro-Electrode Arrays, MEA, 3Brain GmbH) and to maintain sustained electrophysiological activity.

As a second step, we investigated the electrophysiological activity of the developed experimental model by evaluating its capabilities to exhibit spontaneous activity and respond to electrical stimulation. We subsequently computed its spontaneous dynamical richness and perturbational complexity, according to recent metrics also derived from human studies^19,33–36^. Notably, the obtained values approached those found for *in vivo* experimental models while differing from those computed in simpler 2D cultures. Moreover, by increasing the number of co-cultured neurospheroids, i.e., generating assembloids^30,31^, both dynamical richness and complexity increased, suggesting the scalability of our introduced model. Finally, heterogeneity contributed to functional aspects of the network, such as nuanced activity patterns, dynamical variability, and modular coordination. In this context, inhibition both synchronized activity and supported flexible network states, consistent with the literature^37,38^.

In conclusion, our findings indicate that three-dimensionality and modularity are critical for replicating the complexity of *in vivo* brain activity, both in spontaneous and evoked conditions, while cellular heterogeneity acts as a key modulator of network dynamics, enabling more variable activity patterns and stereotyped functional responses. Advancing the development of more sophisticated heterogeneous cultures will be particularly important when evaluating these simplified models against pathological neurospheroids derived from patients with genetic mutations, possibly providing deeper insights into disease mechanisms.

## Methods

### Human induced pluripotent stem cells cultures

We received both Ngn2-positive and Ascl1-positive hiPSCs lines in frozen vials, kindly provided by Prof. Nadif Kasri (Radboud University Medical Centre, the Netherlands). The two hiPSCs lines were previously characterized and genetically modified to generate homogeneous populations of excitatory and inhibitory neurons thanks to the forced expression of the transcription factors neurogenin-2 (Ngn2) and Achaete-scute homolog 1 (Ascl1)^23^. Both lines were generated from fibroblasts. Control line 1 (C1, healthy 30-years-old female) was reprogrammed via episomal reprogramming (Coriell Institute for medical research, GM25256). Control line 2 (C2, healthy 51-years-old male) was reprogrammed via a non-integrating Sendai virus (KULSTEM iPSC core facility Leuven, Belgium, KSF-16-025). The thawed hiPSCs cultures were resuspended into E8Flex medium (Thermo Fisher Scientific) supplemented with E8 supplements (2%, Thermo Fisher Scientific), penicillin/streptomycin (1%, Thermo Fisher Scientific), G418 (50 μg/ml, Sigma-Aldrich) and puromycin (0.5 μg/ml, Sigma-Aldrich). Glutamatergic neurons with cortical-like features were derived from C1 by overexpressing mouse neuronal determinant Ngn2 upon doxycycline (4 μg/ml, Sigma-Aldrich) treatment^67^. GABAergic neurons were derived from C2 by overexpressing mouse neuronal determinant Ascl1 (Addgene, 97329) upon doxycycline treatment and supplementation with forskolin (10 μM, Sigma Aldrich)^23^. The hiPSCs can be considered differentiated into neurons after approximately 3 weeks of doxycycline and forskolin treatment^67,69^. The cultures were maintained in the incubator at stable condition (37°C, 5.5% CO_2_, 95% humidity atmosphere). The medium was refreshed at 50% every two days.

### Rat astrocytes culture

Astrocytes were obtained from wild-type prenatal embryonic rats and were thawed and maintained in high glucose DMEM supplemented with penicillin/streptomycin (1%, Thermo Fisher Scientific), stable L-Glutamine (1%, GlutaMax 100x, Thermo Fisher Scientific) and Fetal Bovine Serum (FBS, 15%, Thermo Fisher Scientific). At DIV 0, astrocytes were detached with Trypsin-EDTA (0.05%, Thermo Fisher Scientific) and were mixed with the human-derived neurons (Fig. 1b) in a neurons:astrocytes ratio equal to 70:30.

**Figure 1:**
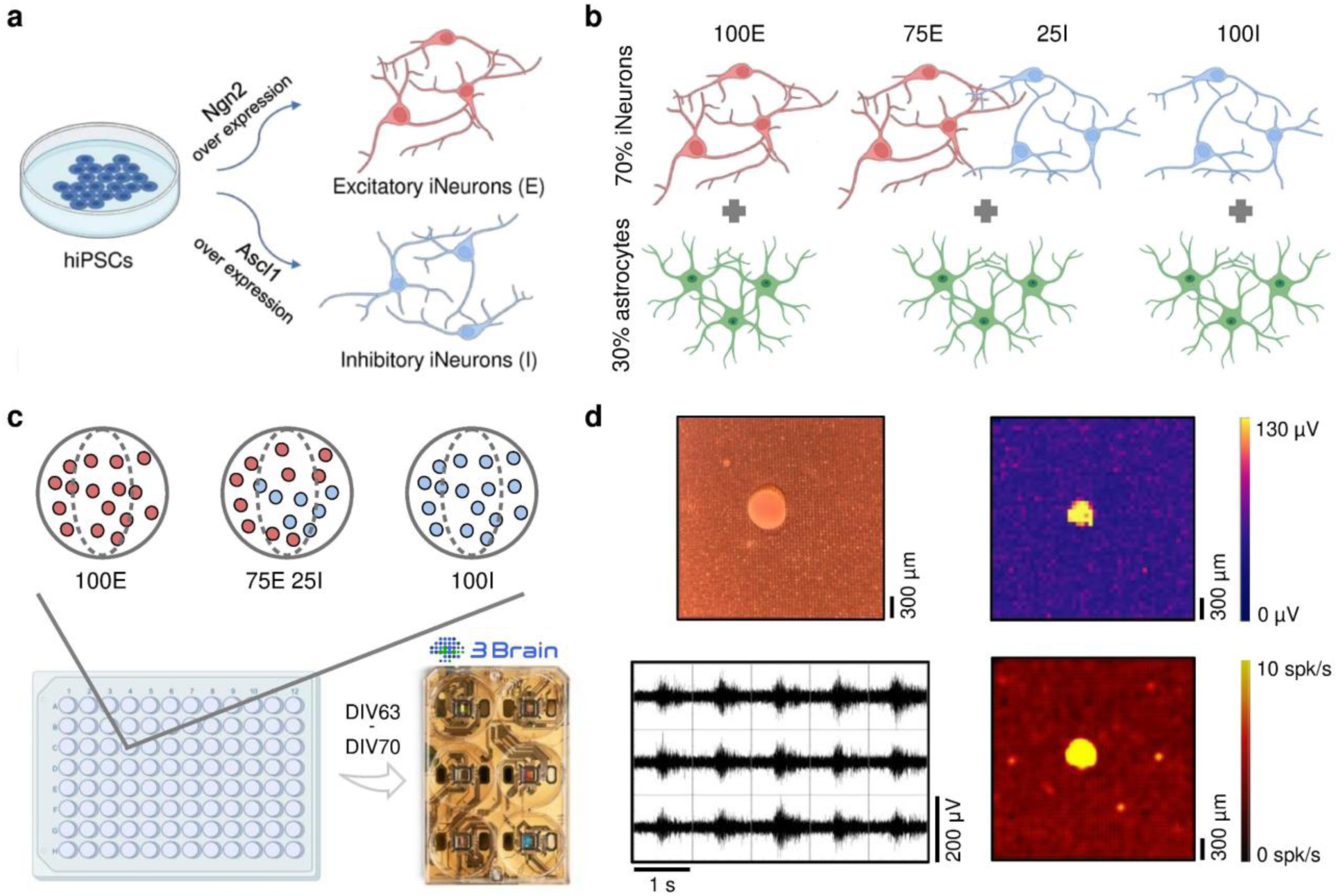
Functional neurospheroids generation. **a)** Glutamatergic (E, in red) and GABAergic (I, in blue) neurons were derived from human induced pluripotent stem cells (hiPSCs) through the overexpression of Ngn2 and Ascl1 transcriptional factors, respectively. **b)** The neurospheroids included 30% of rat astrocytes (in green) to favour the growth and the maturation of the neurons. The three experimental configurations included homogeneous excitatory (100E), heterogeneous (75E25I), and homogeneous inhibitory (100I) neurospheroids. **c)** Neurospheroids were generated by plating the cells in low-adhesion 96-wells to favour the self-organization into a three-dimensional structure and moved on high-density devices (3Brain GmbH) between DIV 63 and DIV 70 to record the electrophysiological activity. **d)** On top left: representative image of a neurospheroid plated on the active area of the high-density MEA; on top right: colour map in which each pixel represents one channel depicting the amplitude (in μV) of the electrophysiological activity of the same representative spheroid; on bottom left: extracellular 1-second signal trace of the same 3D network in which each box represents one channel; on bottom right: firing map of the same spheroid in which each pixel represents the mean firing rate of the channel.

### Neuronal cultures maintenance

During the first week, the neuronal cultures were maintained in Neurobasal medium (Thermo Fisher Scientific) supplemented with B27 supplements (2%, Thermo Fisher Scientific), penicillin/streptomycin (1%, Thermo Fisher Scientific), GlutaMax (1%, Thermo Fisher Scientific), human Brain-Derived Neurotrophic Factor (BDNF, 10 ng/ml, Sigma-Aldrich), human Neurotrophin-3 (NT-3, 10 ng/ml, Sigma-Aldrich), doxycycline (4 μg/ml, Sigma-Aldrich) and forskolin (4 μg/ml, Sigma-Aldrich). After 7 DIV, FBS (2%, Thermo Fisher Scientific) was added to the aforementioned supplemented medium to support astrocytes. After 14 DIV, doxycycline and forskolin were removed from the medium. The neuronal cultures were stably maintained in the incubator (same conditions specified in the previous paragraph). The medium was refreshed at 50% each two days. No exclusion criteria were defined, and all spheroids were included in the analysis. The work has been reported in line with the ARRIVE guidelines 2.0.

### High-density MEAs and electrophysiological recordings

Three days before the electrophysiological recordings, the 3D neuronal cultures were moved from the low-adhesion 96-wells to commercial (3Brain GmbH) CorePlate 6-wells devices or Accura HD-MEA (Supplementary Fig. 5). Indeed, empirical testing indicated that this 3-day settling period prior to MEA recordings provides an optimal balance between stable spheroids adhesion and preservation of their three-dimensional structure. Besides single neurospheroids, we also plated assembloids^30,31^, i.e., two neurospheroids were moved on the same device and put in contact together (Supplementary Fig. 3a, c). The complete dataset of assembloids - utilized for the evaluation of the dynamical richness - is reported in Supplementary Fig. 3b. The CorePlate devices integrated 2304 recording electrodes characterized by 60 μm in pitch and 25 μm in electrodes’ size, arranged in a 2.9 × 2.9 mm^2^ (48 x 48 electrodes) grid, while the Accura HD devices were characterized by a 3.8 × 3.8 mm^2^ (64 x 64 electrodes) grid with 21 μm electrodes and 60 μm pitch. Before moving the neurospheroids, the MEAs were sterilized with 70% ethanol (1h) and UV-lights (45 min) and coated overnight with a mixture of human laminin (20 μg/μl, BioLamina) and poly-L-ornithine (50 μg/μl, Sigma-Aldrich)^22^. Concerning the electrophysiological recordings, after a 10-min period of adaptation out of the incubator, the spontaneous neuronal activity was recorded for 10 minutes (10 kHz sampling rate) in stable condition using a thermostat and a gas mixture cylinder (37°C, 5% CO_2_). The presented results were collected from 4 different batches with 10, 14, 6, and 6 samples each. The recordings were performed once for each neurospheroid between DIV 63 and DIV 70 with the HyperCAM Alpha (with CorePlate devices) or BioCAM DupleX (with Accura devices) systems (3Brain GmbH). The final dataset includes 12 spheroids for 100E configuration, 14 samples for 75E25I condition, and 10 cultures for 100I.

### Electrical stimulation protocol and data analysis

To perform the electrical stimulation ^52^, we tested 20 biphasic (positive-then-negative) pulses characterized by ±25 μA in amplitude, 200 μs in duration, and emitted at 0.1 Hz. Initially, we recorded the spontaneous activity for 10 minutes (see above). During this phase, we chose four electrodes that showed sustained spiking and bursting activity for each neurospheroid, indicating that they were connected to the rest of the network and contributed to the collective activity^70^. The chosen electrodes were subsequently used as stimulation sites. Hence, we stimulated the neuronal networks with the above-mentioned stimulation train from the chosen sites simultaneously, and we recorded the evoked activity for 3.20 minutes. The final dataset for this investigation included 12 100E_30k_ and 10 75E25I_30k_ neurospheroids. The post-stimulus time histograms (PSTHs) were calculated by considering a 1.2 s time window after the stimulus emission. In particular, we divided each time window into 4 ms bins and counted the number of spikes occurring in each bin. We discarded the PSTHs characterized by an area (i.e., the total number of evoked spikes) lower than 2 since these stimuli were considered unable to evoke a response. We computed the percentage of effective stimuli able to evoke a response over a total of 20 stimuli for each stimulation site. Finally, to evaluate the responsiveness of the neuronal network, we calculated the area of the PSTH and the latency, representing the time between the emission of the stimulus and the first evoked spike.

### MEA data analysis

Off-line data analysis was performed using BrainWave software (3Brain GmbH), in-house code developed in MATLAB (The MathWorks, Natick, MA, USA) and SpyCode^71^ to obtain the features describing the spontaneous network activity. Briefly, the spike detection was performed using the precision time spike detection (PTSD) algorithm^72^. The noise threshold for individual spike detection was set at 6 times the standard deviation of the baseline noise, while the peak lifetime period — associated to the duration of the spike— and the refractory period — the minimum time elapsed between consecutive spikes — were set to 2 ms and 1 ms, respectively. Each channel was considered active whether its firing rate was higher than 0.1 spikes/s (spk/s)^73^. The mean firing rate (MFR, i.e., the number of spikes in the unit time) for each 3D culture was computed by averaging the firing rates of each active channel. Moreover, the number of active electrodes (AE) were calculated. Bursts were detected based on the distribution of logarithmic neuronal inter-spike interval (ISI) and setting a threshold of the ISI (set to 100 ms) and the minimum number of spikes belonging to a burst event (set to 10 spikes). An active channel was considered as bursting if its bursting rate was greater than 0.4 bursts/minute^73^. From the burst detection, we extrapolated the mean bursting rate (MBR, i.e., the number of bursts per minute), the percentage of bursting electrodes (BE) with respect to the active channels, the burst duration (BD), the percentage of random spikes (RS, i.e., the number of spikes not belonging to a burst), the peak frequency intra burst (PFIB, defined as the highest instantaneous firing rate observed within a single burst), and the mean frequency intra burst (MFIB, defined as the average firing rate occurring within a burst, calculated as the total number of spikes per burst duration). Finally, the network burst detection was carried out by considering the distribution of the inter-burst-event interval and by setting the minimum number of electrodes participating to the network event equal to the 35%. From the network burst detection, we computed the network bursting rate (NBR, i.e., the number of network events per minute), the network burst duration (NBD), and the bursts within the network bursts (NBs).

### Fragmented network bursts analysis

Since we observed different patterns of network events, some of which characterized by multiple segmentations – referred as fragments – we performed an analysis to detect the amount of fragmented network bursts^49^ and the number of fragments within them. Specifically, by considering the network burst detection described above, if two consecutive network bursts occurred in an interval lower than 2 s, we merged them considering it as a unique fragmented event. Subsequently, we computed the percentage of fragmented NBs – i.e., network events characterized by multiple segmentations occurring in a restricted interval of time – with respect to the total number of network events. Finally, we computed the number of fragments within each fragmented network burst.

### Dynamical richness analysis

To compute the dynamical richness of the networks, we followed the pipeline presented in ref. ^19,36^. Briefly, we evaluated two different contributes: one related to the variability in the pattern of the spiking activity (θ_CC_), and a second one related to the variability in the size of the global network activity (θ_GNA_). θ_CC_ was computed evaluating the pairwise functional Pearson correlation (r_ij_) between the instantaneous firing rates (bin size 100 ms) of two electrodes i and j, from which θ_CC_ was calculated as follows:

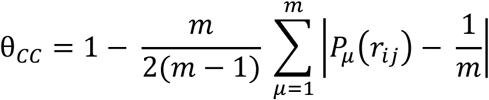

Where m = 20 is the number of bins of the distribution 𝑃_𝜇_(𝑟_𝑖𝑗_).

θ_GNA_ was computed by considering the cumulative firing activity and evaluating the fraction of active electrodes (Γ_t_) within intervals of 0.5 s. Episodes with fewer than 20% of active electrodes were excluded to filter out sporadic, random activity. θ_GNA_ was then calculated as follows:

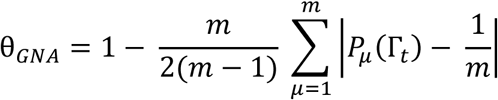

Where m = 20 is the number of bins of the distribution 𝑃_𝜇_(Γ_𝑡_).

Finally, the dynamical richness θ of each network was computed as θ = θ_CC_ · θ_GNA_ . We evaluated both 2D (dataset from ^27^ and ^28^) and 3D neuronal networks, as well as assembloids configuration, that is, two neurospheroids put in contact together on the same device, which included modularity within our model. The final dataset included 50 2D neuronal cultures (27 100E and 23 75E25I), 26 neurospheroids (12 100E and 14 75E25I), and 14 assembloids (Supplementary Fig. 3a).

### Perturbational Complexity Index (PCI) analysis

To compute the perturbation complex index (PCI), we followed the methodology presented in ref. ^33^. First, we discarded the inactive electrodes with a mean firing rate below 0.1 spk/s. For each active electrode, post-stimulus activity within a 1.2 s window was divided into temporal bins of 4 ms. Then we compute the mean response across stimuli, generating an average response matrix. From this, we extracted a binary matrix SS, where values represented significant electrode activations after the emission of the stimuli. The complexity of the SS matrix was quantified using the Lempel-Ziv complexity algorithm. To ensure comparability across conditions, the PCI was defined as the Lempel–Ziv complexity measure normalized between [0,1]. This normalization was performed by dividing the complexity value by the theoretical maximum complexity for a sequence of the same length and entropy. Finally, PCI was defined as follows:

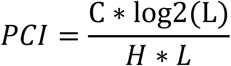

Where C was the Lempel-Ziv complexity, L was the number of elements in the SS matrix and H was the source entropy, defined as:

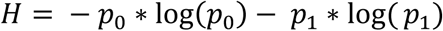

Where 𝑝_0_ and 𝑝_1_were the fraction of 0 and 1 in the SS matrix, respectively. We evaluated the PCI of 2D cultures (dataset from ^22^), 3D neuronal networks, and *in vivo* trials (see below). The final dataset included 12 2D neuronal cultures (6 100E and 6 75E25I), 22 neurospheroids (12 100E and 10 75E25I), and 7 animals obtained from *in vivo* recordings.

### *In vivo* recordings

The *in vivo* data analyzed in this paper are part of a larger dataset acquired in the framework of another study by one of the authors, whose details are reported in ref. ^58,74^. Briefly, the dataset is composed by 7 healthy male Long-Evans rats (weight: 300-400 g; age: 4-5 months; Charles River Laboratories, Calco, LC, Italy). The Italian Ministry of Health and Animal Care (Italy: authorization n. 509/2020-PR) approved all experiments. Recording was done under anesthetized conditions, according to a protocol already published^75^. Anesthesia was maintained throughout the entire procedure with repeated bolus injections of ketamine (10-100 mg/kg/hr intraparenchymal or intramuscular) as needed. Burr holes (3 mm diameter) exposing the premotor cortex (i.e., Rostral Forelimb Area, RFA) and the somatosensory cortex (S1) were made at +3.5, +2.5 and –1.25, +4.25 AP, ML. The cortical activity was recorded in RFA using a four-shank, sixteen-contact site electrode, placed at a depth of 1500 µm (1-1.5 MΩ impedance, A4x4-3mm-100-125-703-A16, NeuroNexus). The stimulation was applied through a single contact on a four-shank, sixteen-contact electrode with an impedance of ∼ 200 kΩ (A4x4-3mm-100-125-177-A16, NeuroNexus). In all the experiments, the stimulation pulse was designed to be a single squared 60 μA biphasic, cathodal-leading pulse (200 μs positive, 200 μs negative). The experimental recordings used in this study consist of 20 minutes of spontaneous activity followed by a few minutes of electrical stimulation (i.e., delivery of 30 stimuli emitted at 0.2 Hz).

### Morphometric analysis

To evaluate the morphometrical characteristics of the neurospheroids, images were acquired from 3D cultures specially designed for this analysis and maintained in the low-adhesion 96-wells. All samples were collected at DIV 63 using an inverted microscope (Olympus IX51, Olympus Corporation) with a 10X objective. Image acquisition settings were standardized across all experimental configurations to compare the images. A total of 103 neurospheroids were acquired, comprising 18 100E_10k_, 17 100E_20k_, 43 100E_30k_, 16 75E25I_30k_ and 9 100I_30k_ neurospheroids. The images were analysed using ImageJ software. Since neurospheroids were not perfect circles, we approximated their shapes to ellipsoid. For each spheroid we measured the length of the major and the minor axes. Subsequently, we averaged the abovementioned values to determine the approximation of the spheroid’s diameter. From the latter, we estimated the perimeters and the area of each spheroid. The circularity - representing how closely the shape of the spheroids approximated a perfect circle - was then calculated as follows:

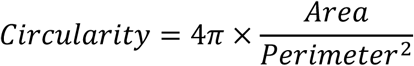

Circularity values approaching 1 indicate a perfect circle, while values closer to 0 represent an increasingly irregular shape.

### Immunocytochemistry

At DIV 70, the neurospheroids were fixed with 4% paraformaldehyde for 15 minutes at room temperature (RT) and permeabilized with 0.2% TritonX-100 for 15 minutes. Subsequently, the samples were exposed to a Blocking Buffer Solution (BBS) consisting of 0.5% bovine fetal serum and 0.3% bovine serum albumin in PBS for 45 minutes at room temperature. The primary antibodies were diluted in BBS and incubated at room temperature for 4 hours. The following labelling were performed: 1) DAPI, to evaluate nuclear morphology.; 2) Anti-GABA + NeuN, to evaluate different compositions of the heterogeneous cultures; 3) MAP2 + GFAP, to evaluate the cellular composition inside the neuronal networks at DIV70, distinguishing neuronal and astrocytic populations. The complete description and dilution of primary antibodies is reported in Supplementary Fig. 2a. Depending on the primary antibodies utilized, a combination of the following secondary antibodies was used: Alexa Fluor 546 goat anti-guinea pig (1:1000, Invitrogen), Alexa Fluor 546 goat anti-rabbit (1:1000, Invitrogen), Alexa Fluor 488 goat anti-rabbit (1:700, Invitrogen), Alexa Fluor 488 goat anti-mouse (1:700, Invitrogen). All the samples were treated with DAPI (1:5000, Sigma-Aldrich) for 10 minutes at RT to label cellular nuclei.

Confocal imaging was acquired on a Nikon Eclipse upright microscope coupled with 20× Nikon objective, 0.80 NA (Nikon CFI Plan Apochromat Lambda D). The image-processing package Fiji ^76^ was used for further analysis of the acquired data. To distinguish live and dead neuronal cells, nuclear fragmentation and nuclear size were evaluated in cells labelled with DAPI. Images were first converted to 16-bit, a threshold was applied, and the images were binarized. The watershed algorithm was then used to separate clusters of cells into single cells. Finally, the ‘*Analyze Particles*’ function was applied to select nuclear areas corresponding to the dimensions of live and dead cells. An average nuclei size was calculated and used as the reference for quantification^77^. Nuclear fragmentation and nuclei reduced by ≥ 50% in size compared with live controls were defined as indicators of cell death. For each condition, three samples were considered. Four fields 98x98 μm^2^ (Supplementary Fig. 2d) and 3 fields 140x140 μm^2^ with a 512x512 resolution and Z-step = 4 μm, for a total of 25 layers were acquired for each sample. Images of neurospheroids with the labelling 2) and 3) were acquired of the whole field with a 2048x2048 resolution with Z-step 2 μm for around 150 layers.

### AFM acquisition and data analysis

To evaluate the mechanical properties of single neuronal cells and neuronal spheroids, force spectroscopy measurements were conducted using an atomic force microscope (AFM, JPK NanoWizard 4, Bruker Corp., Berlin, Germany) coupled with an upright optical microscope (Zeiss Axio Zoom V16). The optical access to the AFM probe and sample facilitated the identification of the region of interest for acquisition. For all measurements, the same rectangular cantilever with a pre-calibrated spring constant of 0.182 N/m and a spherical tip with a 5 µm radius (SAA-SPH-5UM, Bruker Corp., Berlin, Germany) were used. Before starting the measurements, the deflection sensitivity was assessed by performing force-distance curves on a glass slide in the same culture medium used with the cells. Force spectroscopy measurements were performed on each sample adhering to plastic petri dishes placed on a heated stage (37°C). Maps of force-distance curves with a pixel resolution of 5 µm were acquired utilizing the Smart Mapping software (Bruker Corp., Berlin, Germany) which allowed to arbitrary define scan areas and combine both piezo scanner and step motor movements to achieve large scan sizes and evaluate tall sample features. The force set-point was set to 3 nN and the vertical movement speed to 16 µm/s. *<u>2D cultures</u>:* bidimensional neuronal cultures composed of exclusively excitatory or inhibitory neurons were evaluated by AFM. Maps containing either individual cells or small clusters were acquired. We minimized the contribution of the substrate by discarding substrate-related curves during the post-processing. The final dataset included 4307 and 4383 curves for the excitatory and inhibitory neurons, respectively. *<u>3D cultures:</u>* three-dimensional spheroids of the considered configurations, i.e., 100E, 75E25I, and 100I, were evaluated by AFM. The final dataset included 5699, 2124 and 615 curves for the 100E, 75E25I, and 100I configurations, respectively.

AFM force-distance curves were processed using the JPK Data Processing software (Bruker, Berlin, Germany) to extract force-indentation data and calculate the Young’s modulus (E) according to the Hertz/Sneddon model for spherical indentation^78^. Specifically, the relationship between the loading force (F) and the penetration depth (δ) was defined by the following equation:

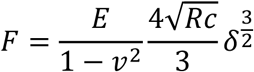

Where E was the elastic modulus, 𝑅𝑐 was the radius of the spherical indenter (5 µm), δ was the indentation depth, and 𝑣 was the Poisson’s ratio (set to 0.5 for incompressible materials).

### Statistical analysis

Statistical analyses were performed using MATLAB (The MathWorks, Natick, MA, USA). We evaluated the normal distribution of the data using the Kolmogorov–Smirnov normality test. For not normally distributed data, we performed a non-parametric Mann-Whitney (for pairs comparison) or Kruskal–Wallis with Dunn’s correction test (for multiple comparison). For normally distributed data, we performed an unpaired t-test (for pairs comparison) or One-way ANOVA with Tukey’s correction (for multiple comparison). To establish statistically significant differences, p-values < 0.05 were considered significant.

## Results

### Engineered neurospheroids generation

To realize engineered functional neurospheroids, we mixed 70% of human-derived excitatory (E) and inhibitory (I) neurons (iNs, Fig. 1a) at different E/I ratios with 30% of rat astrocytes (A, Fig. 1b), following an experimental pipeline already adopted in previous studies^27^. 3·10^4^ cells were plated in low-adhesion 96-wells (Fig. 1c) to favour the self-organization into 3D structures^39,40^. We created homogeneous excitatory neurospheroids in which exclusively glutamatergic neurons were included (referred to as ‘100E’ configuration). The heterogeneous configuration consisted of 75% of glutamatergic neurons and 25% of GABAergic ones (referred to as ‘75E25I’ configuration). Finally, the homogeneous inhibitory neurospheroids (referred to as ‘100I’ configuration) were created by plating only hiPSCs-derived GABAergic neurons. Moreover, to evaluate the effects of the 3D culture dimensionality, we created homogeneous excitatory neurospheroids (100E) with 10^4^, 2·10^4^, and 3·10^4^ cells. Three days before the electrophysiological recordings, the 3D neuronal cultures were moved from the low-adhesion 96-wells to high-density MEAs (2304 or 4096 electrodes, 3Brain GmbH, Fig. 1c), which allowed to record the extracellular signal traces, thus providing high-resolution recordings of spiking activity across the 2D layer interfacing with the MEA (Fig. 1d). The neurospheroids successfully adhered to the MEAs and were fully functional along the entire surface in contact with the devices. Moreover, the 3D cultures exhibited an organized activity characterized by the simultaneous activation of several channels, proving a synchronicity of the network (Fig. 1d).

### Morphomechanical characterization

With the aim of obtaining a fully controlled *in vitro* neurospheroid system, we first evaluated different compositions in terms of heterogeneity and number of cells. In particular, we focused on the morphological variations of the neurospheroids with different amounts of cells, i.e., 3D excitatory neuronal cultures (100E) composed of 10k, 20k, or 30k cells. By qualitatively observing the optical images of representative neurospheroids (Fig. 2a), the amount of cells within the cultures was reflected in the dimension of the corresponding spheroids. Indeed, the diameter of the 3D structures showed increasing values proportional to the number of cells (Fig. 2b). Clearly, the variations were equally marked considering the area of the neurospheroids, which demonstrated higher average values when 30k cells were included in the 3D cultures (Fig. 2b). Finally, both 20k and – especially – 10k configurations showed lower circularity with respect to the 30k cultures, suggesting that the structure of these neurospheroids was characterized by a more irregular shape. The changes in the dimensionality were reflected in the firing pattern, as 30k spheroids showed a sharp increase in the mean firing rate (MFR), while 10k and 20k spheroids revealed comparable MFR values (Supplementary Fig. 1a).

**Figure 2:**
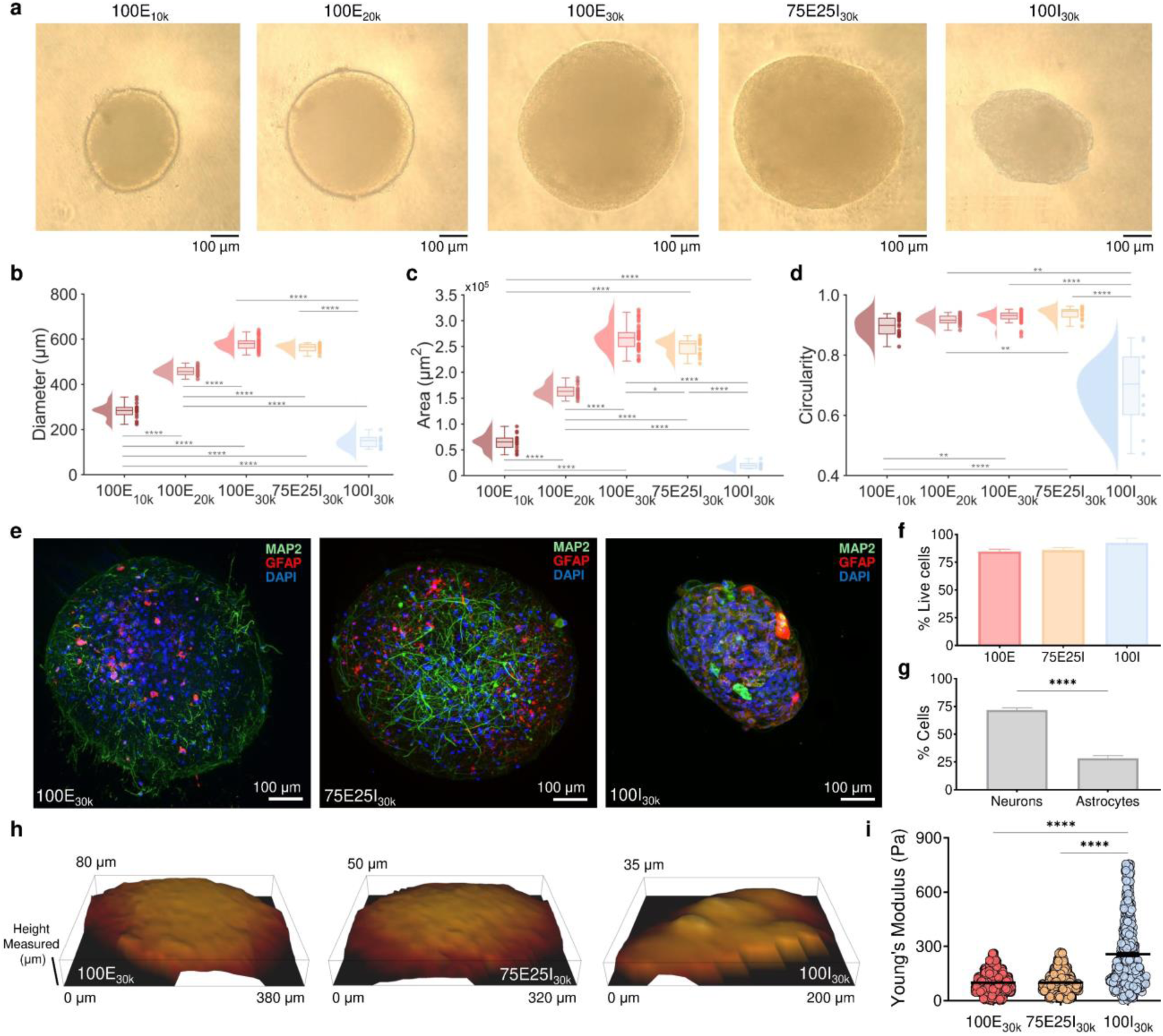
Morphomechanical characterization. **a)** From left to right: representative images of 100E_10k_, 100E_20k_, 100E_30k_, 75E25I_30k_, and 100I_30k_ neurospheroids. **b-d)** Box plots with violins (on the left) and scatters (on the right) of **b)** average diameter; **c)** area; and **d)** circularity of the 3D cultures for each condition (N_100E_10k_ = 18; N_100E_20k_ = 17; N_100E_30k_ = 43; N_75E25I_30k_ = 16; N_100I_30k_ = 9). In the box plots, data are represented with the percentile 25-75 (box), the standard deviation (whiskers), and the median (line) values (One-way ANOVA or Kruskal-Wallis’ test, * refers to p < 0.05. ** to p < 0.01, *** to p < 0.001, **** to p < 0.0001). **e)** Immunofluorescence images of 100E, 75E25I and 100I (from left to right) neurospheroids labelled with MAP2 (green), GFAP (red), and DAPI (blue). **f)** Box plot of the amount of live cells within neurospheroids for each configuration. Data are represented with percentile 25-75 (box), median (line) and percentile 10-90 (whiskers). N_100E_ = 103; N_75E25I_ = 83; N_100I_ = 16 (Kruskal-Wallis’ test with significance’s level p < 0.05). **g)** Box plot of the amount of neurons and astrocytes within neurospheroids. Data are represented with percentile 25-75 (box), median (line) and percentile 10-90 (whiskers). N = 177 (Mann-Whitney test, **** refers to p < 0.0001). **h)** Topological three-dimensional reconstruction of three representative spheroids (one for each configuration) obtained from Atomic Force Microscopy acquisition. **i)** Scatter plots of Young’s Modulus for each configuration (N_100E_30k_ = 5699; N_75E25I_30k_ = 2124; N_100I_30k_ = 615). Data are represented with the mean (horizontal line) and the standard error of the mean (whiskers) (Kruskal-Wallis’ test, **** refers to p < 0.0001).

Regarding the morphometric characterization of the other configurations, i.e., 75E25I_30k_ and 100I_30k_, the heterogeneous neurospheroids with respect to 100E_30k_ showed similar shapes (Fig. 2a) with comparable diameter (Fig. 2b), a slightly lower area (Fig. 2c), and highly comparable circularity (Fig. 2d) with values approaching 1, suggesting almost perfect spherical shape. On the other hand, the 100I_30k_ neurospheroids showed a strongly reduced circularity and dimension with respect to 100E_30k_ and 75E25I_30k_, despite the initial amount of cells included equal to 30k. Nevertheless, all neurospheroids showed a high amount of live cells (over 84%) in every configuration (Fig. 2f, Supplementary Fig. 2b) and a network characterized by neurons (MAP2 positivity, Fig. 2e, and Supplementary Movies 1-3) and astrocytes (GFAP positivity, Fig. 2e, and Supplementary Movies 1-3) with an iNs:A ratio equal to 72:28 (Fig. 2g), highly consistent with the declared nominal values plated at Day *In Vitro* (DIV) 0. Differences in the morphology were also reflected in both the topographical images (Fig. 2h, Supplementary Fig. 1b) and the stiffness values (Fig. 2i, Supplementary Fig. 1c) obtained by Atomic Force Microscopy (AFM) measurements. The latter showed higher values of the Young’s module for the 100I_30k_, and comparable values for the configurations in which the excitation was present.

In view of the above, we selected 30k as the target configuration for our neurospheroids and, from this point onwards, we omit the subscript ‘30k’ to indicate the spheroids with 3·10^4^ cells, and we refer to the three different configurations as 100E, 75E25I, and 100I.

### Spontaneous activity of the neurospheroids

The neurospheroids generated through our optimized protocol revealed robust, sustained spontaneous activity, with distinct phenotypes depending on the E/I balance. Upon reaching full maturation (i.e., after DIV 50 upon GABA shift^23^), they exhibited a combination of single-channel spiking and bursting activity as well as synchronized network burst events, as illustrated in Fig. 3. Representative instantaneous firing rate profiles are shown in Fig. 3a, which reports snapshots of spontaneous activity for the different considered configurations, i.e., 100E, 75E25I and 100I. Both 100E and 75E25I cultures showed dynamic patterns referable to bursting and network bursting activity, while 100I neurospheroids exhibited tonic firing patterns lacking synchronized events. Additionally, network bursts in both 100E and 75E25I cultures were characterized by a fragmented shape (Fig. 3a, b), particularly prominent in the 100E configuration. By evaluating key features which describe the dynamics of the electrophysiological activity, the radar plot of Fig. 3c reveals distinct behaviour across the three configurations. The pure inhibitory configuration (100I) showed a lower number of active electrodes, consistent with their smaller size (cf. Fig. 2). Moreover, both the mean bursting rate (MBR) and network bursting rate (NBR) for the 100I were zero as reflected in the raster plot. This result aligns with the tonic firing activity observed in these neurospheroids, which exhibited only random spiking without synchronized events (Fig. 3c). In contrast, when considering the bursting patterns of the configurations which included the excitatory component (100E and 75E25I; Fig. 3c), no statistical differences were found between the two, despite the substantial difference introduced by the presence of GABA in the 75E25I spheroids (Supplementary Fig. 2c). Finally, given the varying levels of network burst fragmentation observed in the qualitative profiles (Fig. 3a), we conducted a quantitative analysis to assess differences in the network bursting patterns. Consistent with the qualitative findings, all 100E cultures showed fragmented NBs in 20% to 100% of their total network events. In contrast, 4 distinct 75E25I neurospheroids from 3 different batches (indicating that the observation is not batch-dependent) did not display fragmented NBs, suggesting a broader repertoire of network event shapes (Fig. 3d). Considering the number of fragments within the cultures that exhibited such a pattern (Fig. 3e), the 100E configuration exhibited between 2 to 9 fragments, while the 75E25I configuration showed a higher degree of variability, with some cultures displaying more than 10 fragments, indicating a more diverse dynamic repertoire.

**Figure 3:**
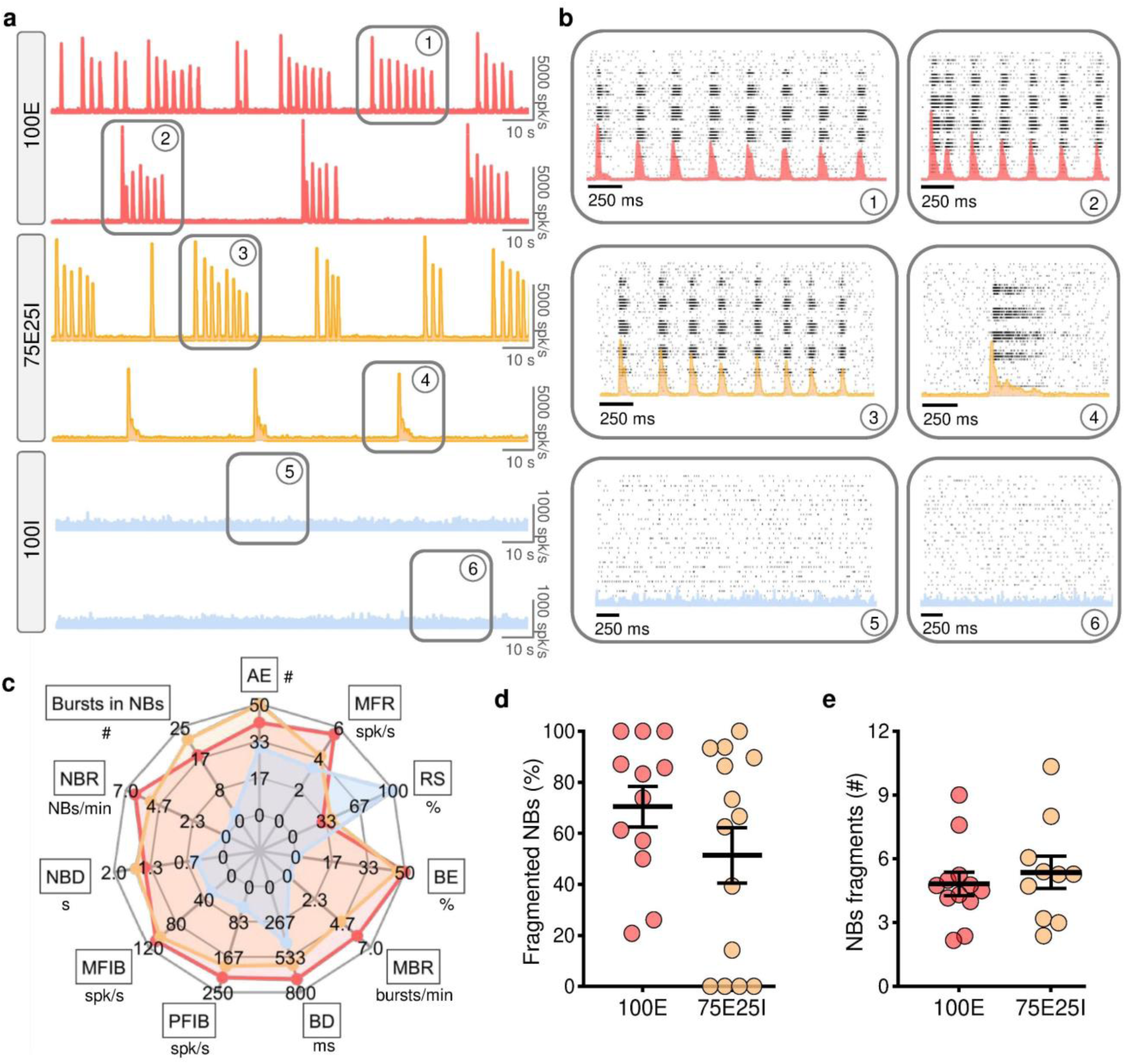
Spontaneous activity characterization of 30k neurospheroids. **a)** Representative cumulative instantaneous firing rate profiles (bin size = 10 ms) of 100E, 75E25I, and 100I neurospheroids (from top to bottom, respectively). **b)** Raster plots zoom with overlapped the instantaneous firing rate profile (bin size = 10 ms) of the electrophysiological activity highlighted with grey boxes in the a) panel. A black dot and a dense black band represent a detected spike and a network burst, respectively. **c)** Radar plot for each configuration (100E in red, 75E25I in orange, 100I in blue) reporting the number of Active Units (AE); Mean Firing Rate (MFR); percentage of Random Spikes (RS); percentage of Bursting Electrodes (BE); Mean Bursting Rate (MBR); Burst Duration (BD); Peak Frequency Intra Burst (PFIB); Mean Frequency Intra Burst (MFIB); Network Burst Duration (NBD); Network Bursting Rate (NBR); and number of bursts within network bursts (NBs). **d-e)** Scatter plots of **d)** percentage of fragmented network bursts with respect to the total amount of network events and **e)** number of fragments within fragmented NBs. In the scatter plots, data are reported with the mean (horizontal line) and the standard error of the mean (whiskers). N_100E_ = 12; N_75E25I_ = 14; N_100I_ = 10.

### Electrical stimulation responsiveness

Our neurospheroids demonstrated also robust and consistent responses to electrical stimulation, with high levels of network recruitment across the cultures. Electrical stimulation of both 100E and 75E25I cultures was able to effectively trigger network burst events, showing a strong correlation between the timing of the stimuli and the evoked activity, as qualitatively depicted in the raster plots of Fig. 4a. The post-stimulus time histogram (PSTH) was characterized by a canonical shape with a straight rising phase and a smooth decay phase in both configurations (Fig. 4b). The area underneath the curve of each channel reflected the signal propagation from the stimulation site across the entire spheroids, with higher values near the stimulation channel and decreasing towards the periphery (Fig. 4c). By quantifying the evoked responses (Fig. 4d), both configurations showed a comparable number of positive responses (about 75% for 100E and 69% for 75E25I), quantitatively confirming a high recruitment of the networks. Moreover, they showed almost immediate responsiveness of the networks with latency values below 50 ms for both configurations (Fig. 4e). The normalized area of the PSTH highlighted a trend towards lower values for the 75E25I, although not statistically significant (Fig. 4f). Despite the response to the electrical stimulation did not highlight strong differences between the two configurations, we assessed the existence of morphological differences within the neurospheroids by labelling GABA neurotransmitter to confirm the presence of inhibitory neurons within the model (Supplementary Fig. 2c). Indeed, the 75E25I cultures were positive for GABA, showing a clear and strong immunoreactivity, contrarily to the 100E spheroids (Supplementary Fig. 2c). This evaluation proved that differences in the internal neural composition of the 3D models were not reflected in the responsiveness to the electrical stimulation, which successfully evoked a comparable response in the neurospheroids.

**Figure 4:**
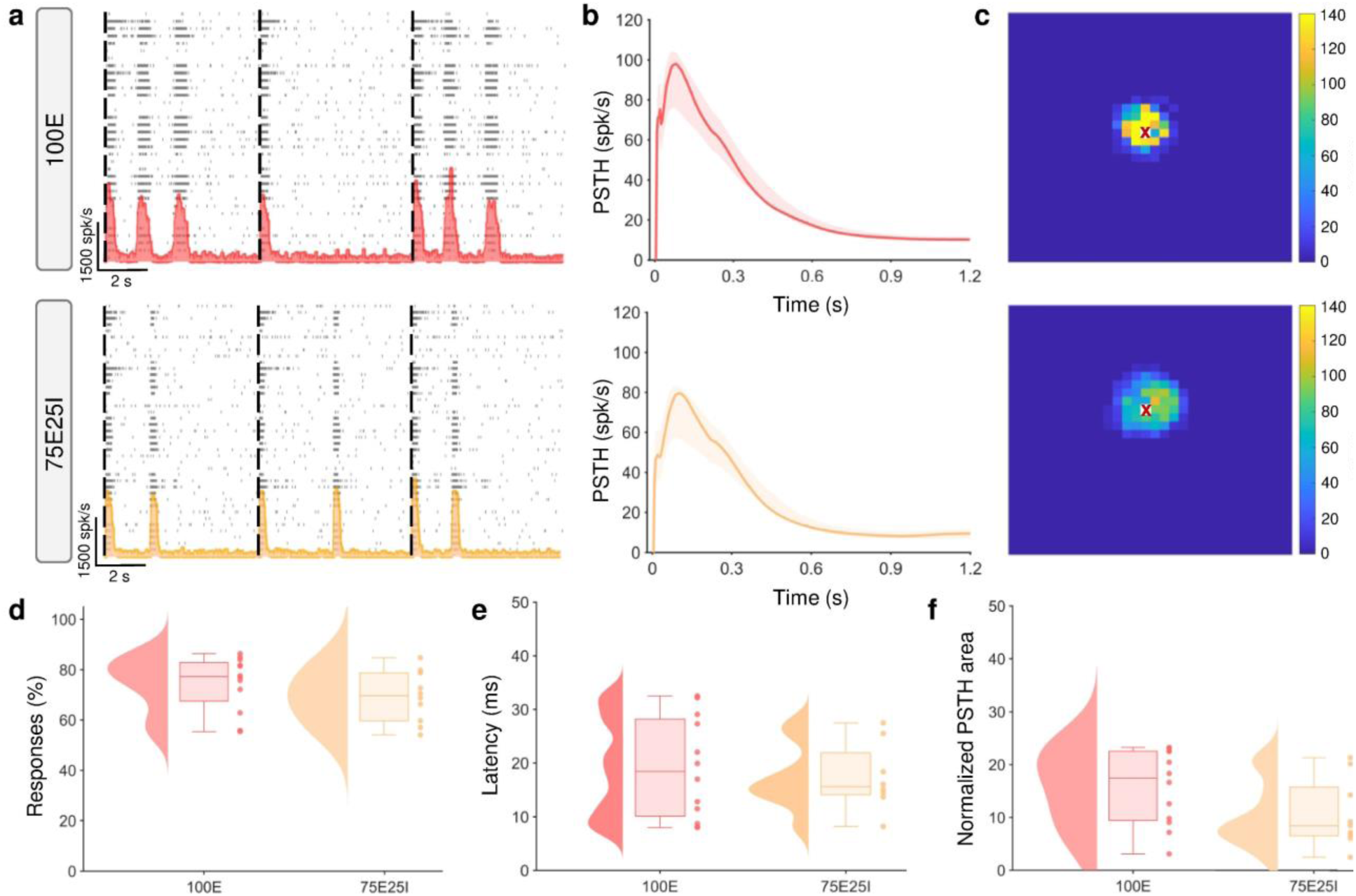
Electrically evoked activity characterization of 30k neurospheroids. **a)** Raster plots of representative 100E (on top) and 75E25I (on bottom) spheroids with overlapped the cumulative instantaneous firing rate profiles (bin size = 10 ms). A black dot and a dense black band represent a detected spike and a network burst, respectively. Black dashed lines overlapped to the raster plots represent the timing of emission of the stimuli (0.1 Hz). **b)** PSTH average profiles (thick lines) of 100E (on top) and 75E25I (on bottom) configurations with 80% confidence intervals (shadows). **c)** Colour maps representing the area underneath the PSTH curves for each electrode of 100E (on top) and 75E25I (on bottom) representative spheroids. Red crosses represent the stimulation site. **d-f)** Box plots with violins (on the left) and scatters (on the right) for each configuration of **d)** percentage of positive responses; **e)** Latency; **f)** Normalized PSTH area. In the box plots, data are represented with the percentile 25-75 (box), the standard deviation (whiskers), and the median (line). N_100E_ = 12; N_75E25I_ = 10; unpaired t-test

### Dynamical richness and perturbational complexity

To assess whether our neurospheroids and assembloids – a modular model where two spheroids are in contact (see Methods, Supplementary Fig. 3a) – are able to capture the rich dynamics and complexity of *in vivo* brains and differ from 2D networks, we quantified their dynamical richness ^19^ and perturbational complexity ^33,34^ (Fig. 5). We first computed the dynamical richness (see Methods) of 2D and 3D neuronal networks. Both 100E (Fig. 5a, Supplementary Fig. 4a, d) and 75E25I (Fig. 5b, Supplementary Fig. 4b, e) neurospheroids showed significantly higher dynamical richness with respect to the 2D models realized with the same neuronal composition (i.e., homogenous excitatory and heterogeneous), proving a richer pattern in the 3D model. To include modularity in our model, we created assembloids in all possible different combinations to have a heterogeneous dataset (Supplementary Fig. 3b). We evaluated changes related exclusively to the model (i.e., 2D, 3D single spheroids, and 3D assembloids) and not to the composition of the cultures (excitation or inhibition). For this reason, we merged all the data of the same model, independently of the cellular configuration (Fig. 5c, Supplementary Fig. 4c, f). Indeed, our models exhibited an increasing dynamical richness from 2D to 3D assembloids, with statistical differences between 2D and 3D models. Notably, 3D assembloids revealed higher values of the dynamical richness’ component related to the functional correlation with respect to the single spheroid model (Supplementary Fig. 4c).

**Figure 5:**
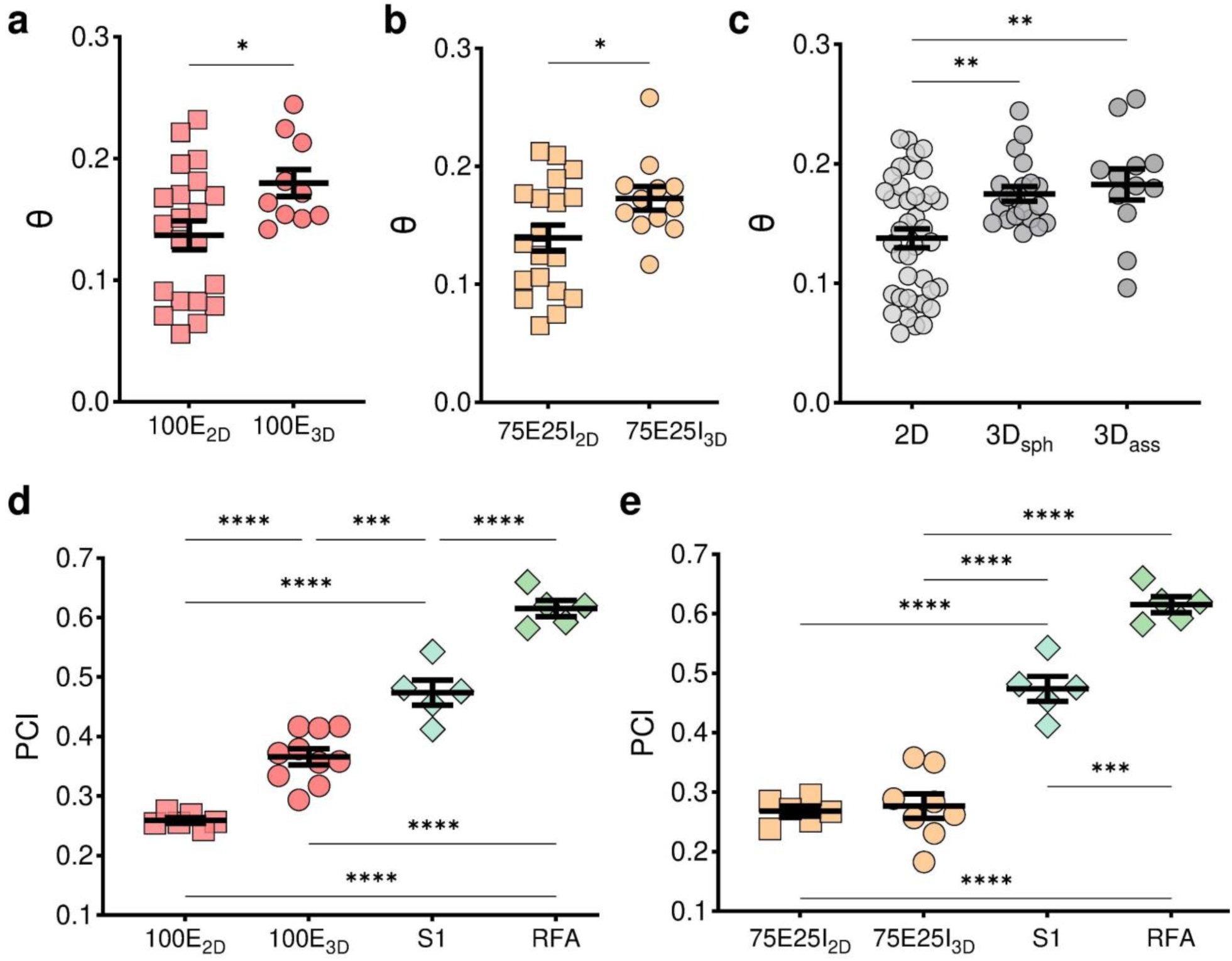
Dynamical richness (θ) and perturbational complexity index (PCI). **a)** Dynamical richness of 2D and 3D 100E neuronal networks (N_100E_2D_ = 27; N_100E_3D_ = 12). **b)** Dynamical richness of 2D and 3D 75E25I neuronal networks (N_75E25I_2D_ = 23; N_75E25I_3D_ = 14). **c)** Dynamical richness of 2D, 3D spheroids and 3D assembloids models, independently from the culture composition (N_2D_ = 50; N_3D_sph_ = 26; N_3D_ass_ = 14). **d)** Perturbational complexity index of 2D and 3D 100E neuronal networks and *in vivo* data of S1 and RFA stimulation (N_100E_2D_ = 6; N_100E_3D_ = 12; N_S1_ = N_RFA_ = 7). **e)** PCI of 2D and 3D 75E25I neuronal networks and *in vivo* data of S1 and RFA stimulation (N_75E25I_2D_ = 6; N_75E25I_3D_ = 10; N_S1_ = N_RFA_ = 7). In the scatter plots, data are represented with the mean (horizontal line) and the standard error of the mean (whiskers). (* refers to p < 0.05, ** to p < 0.01, *** to p < 0.001, **** to p < 0.0001; a) and b) Unpaired t-test; c), d), and e) One-way Anova test).

The analysis of the Perturbational Complexity Index (PCI) was performed comparing 2D, 3D and *in vivo* models under stimulation regimen (see Methods). The PCI of the *in vivo* data reasonably showed higher values than both 100E and 75E25I models (2D and 3D), especially in the case of the premotor cortex (i.e. Rostral Forelimb Area, RFA in Fig. 5d, e). Moreover, the three-dimensional model of the 100E proved to have straight increased values of PCI compared to the same 2D model (Fig. 5d). Conversely, 75E25I 3D model exhibited a very slight and not statistical increase of the PCI with respect to 2D model (Fig. 5e).

## Discussion

In this study, we introduced a simple and reliable engineered human model that represents a crucial first step in developing an *in vitro* system capable of replicating complex *in vivo* features. Specifically, we successfully generated functional neurospheroids that adhered to the surface of high-density MEAs and demonstrated peculiar electrophysiological activity. We effectively controlled uniform-sized neurospheroids, as confirmed by the morphometric analysis, which showed that increases in diameter and area corresponded to a higher MFR. We identified an optimal size, in terms of cells’ number, that ensured a consistent and repeatable circular shape and sustained spiking activity, essential for in-depth studies of network dynamics. Notably, the networks exhibited a high percentage of viable cells, suggesting that the necrotic core of the neurospheroids had been minimized. Additionally, we successfully maintained a neuron-to-astrocyte ratio of about 70:30, and immunofluorescence images revealed a high presence of GABA in the 75E25I neurospheroids, further highlighting the differences in cellular composition between the homogeneous and heterogeneous configurations. Importantly, previous investigations have established that the 75E25I ratio is stably preserved over extended culture periods in 2D systems^27^, strongly suggesting that a comparable equilibrium is likewise sustained in the 3D context. Furthermore, in the same cellular platform, the functional integration of inhibitory neurons has been rigorously validated through MEA recordings performed before and after pharmacological blockade with GABA_A_ antagonists^28^, which elicited alterations in network dynamics. Stiffness analysis conducted with AFM images revealed Young’s Modulus comparable to 3D rat-derived neurospheroids^41^ and higher in the 100I configuration compared to the 100E and 75E25I groups, which displayed similar values. This difference in the morphomechanical properties of the 100I configuration is likely related to the aggregation process of the 3D structure, as the stiffness of inhibitory and excitatory single neurons exhibited comparable values (Supplementary Fig. 1c). To the best of our knowledge, this is the first study to employ AFM to assess the stiffness of individual hiPSC-derived neurons with a defined excitatory or inhibitory phenotype. Indeed, existing literature has predominantly characterized neuronal mechanical properties based on their anatomical area of origin, without distinguishing between excitatory and inhibitory identities^42,43^. Additionally, 100I neurospheroids exhibited a reduced size, resembling the morphometric characteristics of the 10k ones. Since the initial number of cells used to build up the model was the same as in the 100E and 75E25I groups (i.e., 30k cells), this size reduction suggests a loss of cells that were not incorporated into the 3D structure. It is likely that the inhibitory neurons in a pure network were hindered by the absence of excitatory neurons, which are crucial for the development and maturation of GABAergic neurons^44,45^. It is worth noting that we were the first to present human-derived 2D^27^ and 3D (this work) homogeneous inhibitory networks, since - to the best of our knowledge - the literature lacks electrophysiological characterization of such cultures. This underlines the need for further protocol optimization, outlining that the 100I model represents a delicate and challenging configuration to manage, especially when dealing with 3D architecture.

The analysis of electrophysiological activity revealed that the 100I neurospheroids were characterized solely by tonic firing, without any bursting or network bursting events, consistent with findings from purely 2D inhibitory networks^46–48^. In contrast, both the 100E and 75E25I configurations exhibited bursting and network bursting activity, with the appearance of fragmented network bursts^49^. In the 75E25I networks, only a subset displayed these events and exhibited a higher number of fragments within the network bursts; while in the 100E configuration, all neurospheroids exhibited this pattern, highlighting the more diverse dynamics of the heterogeneous configuration. Conversely, no significant differences were observed in the evaluation of electrical stimulation, which elicited a comparable response in both configurations. This demonstrated good recruitment and responsiveness of the entire network in both homogeneous and heterogeneous models, despite the different compositions of the cultures. It is worth highlighting that the heterogeneous 3D networks responded more reliably to electrical stimulation than what was observed in two-dimensional networks^50–52^.

Lastly, we performed the analysis of dynamical richness and PCI. The dynamical richness describes the complexity and diversity of behaviours exhibited by a dynamical system over time^19,36^. We first compared the 2D and 3D networks in both homogeneous and heterogeneous configurations. Three-dimensional networks exhibited a richer activity pattern compared to their 2D counterparts across both configurations. Subsequently, to demonstrate the scalability of our model, we analysed this feature by considering 2D, 3D single spheroids (3D_sph_), and 3D assembloids (3D_ass_), focusing on the model itself rather than on neuronal composition. The richness was further enhanced by increasing the number of spheroids in the 3D_ass_ model, with this effect being mainly driven by the spiking activity component, underlining the relevance of introducing key features such as modularity and three-dimensionality to shape network behaviour. On the other hand, we computed the PCI to describe the complexity of responses to perturbations, quantifying the spatiotemporal diversity of neural activity and serving as an indicator of network integration and differentiation^33^. It is commonly accepted that *in vitro* networks are considered non-awake^53–56^; therefore, to ensure consistency in our comparison, we rationally included *in vivo* data of electrical stimulation performed on anesthetized animals^57,58^. Although the *in vivo* dataset differs from the spheroid recordings in species, preparation, and stimulation protocols, it served as a qualitative reference to contextualize the PCI values. Despite these methodological differences, the comparison provides an approximate indication of how spheroid complexity related to *in vivo* neural dynamics. Indeed, the PCI analysis performed on our networks revealed a gradual and proportional increase in PCI, reaching maximum values with the *in vivo* data. In particular, showing a clear upward trend, the 100E networks exhibited a higher level of complexity, with the 3D networks approaching *in vivo* values. In contrast, the 75E25I networks displayed similar average PCI values in both 2D (PCI = 0.27) and 3D (PCI = 0.28) models. This is likely due to the role of inhibition in regulating activity by synchronizing network events, leading to a more patterned electrical stimulation response and resulting in a less complex response and – consequently – a lower PCI^59–61^.

Supporting our findings, previous studies have systematically compared neuronal dynamics in 2D and 3D *in vitro* networks, highlighting the impact of dimensionality on functional complexity. For instance, Severino et al. reported that 3D rat-derived cultures exhibit network activity regimes rarely observed in 2D, including both highly synchronized and moderately synchronized states^62^. Unlike the stereotyped synchronous bursting typical of 2D networks, 3D networks display more variable and locally modulated synchronization patterns, which more closely resemble *in vivo* dynamics. Similarly, Frega et al. demonstrated that 2D rat networks are dominated by strong synchronous bursting with few random spikes, whereas 3D networks show a significantly higher proportion of random spiking and a reduced bursting rate, again indicating a richer dynamical repertoire^63^. More recently, Winter-Hjelm et al. showed that 2D rat cultures mature into hyperexcitable networks with high mutual information, strong clustering, and elevated bursting rates, but at the expense of segregated information processing^64^. Overall, these findings suggest that while 2D networks tend to develop highly interconnected and stereotyped activity patterns, 3D networks achieve a richer and more flexible dynamical repertoire, reflecting a higher degree of functional complexity.

Despite the innovative approach taken in our study of these human-derived networks, there are a few limitations worth noting. First, while we employed high-density devices, recordings were still planar, measuring only the electrophysiological activity at the base of the neurospheroids. Notably, previously published studies using human-derived experimental models still rely on planar devices for electrophysiological recordings with MEAs^40,65^. Future studies should integrate techniques (which, at the moment, are not commercially available) to assess the activity within the spheroid itself. Additionally, it is important to mention that the spheroids were generated by incorporating rat astrocytes, resulting in mixed-species networks. This choice was driven by the fact that protocols for deriving human astrocytes from hiPSCs are recent, and many aspects of their development remain unclear^66^. In contrast, rat astrocytes are well-characterized, commonly used, and have been previously employed with these specific hiPSCs lines^23,67^; while this introduces inherent interspecies variability, its impact is considered minimal in the context of our network analyses. Indeed, although our approach relies on a heterologous support system, the electrophysiological activity we record originates from human neurons, while rat astrocytes act indirectly by sustaining neuronal and synaptic development^68^. Species-specific differences, such as faster rodent maturation and calcium metabolism, may still shape the circuit dynamics; however, this setup allows us to capture human neuronal function within a supportive environment, which is particularly valuable for translational and personalized medicine applications. Nevertheless, a fully human model is a promising future direction to better assess astrocytic contributions to the network. Moreover, while assembloids offer a promising approach to integrate modularity, this work should be considered a preliminary exploration, providing a useful starting point to capture global trends. Future studies, based on larger dataset, will aim at a more comprehensive analysis of assembloids, including the characterizing composition-specific contributions. Finally, the dynamical richness was computed with relatively coarse bin sizes which may smooth over finer temporal dynamics that could contribute to network complexity, particularly in relation to inhibitory neurons’ activity; and the PCI analysis was performed using a sparse stimulation protocol (four electrodes, low-frequency pulses), which may limit full engagement of underlying network dynamics.

In summary, despite the possibility for further enhancements to our presented model, our comprehensive analysis of richness and complexity provides strong evidence that our engineered three-dimensional model resembles the characteristics of *in vivo* networks and exhibits a functional behaviour that is both rich and complex. This suggests that human-based neurospheroids and assembloids constitute a promising step forward in approximating cortical complexity. In particular, three-dimensionality and modularity were mainly responsible for a significant increase in richness compared to the traditional 2D model, across both homogeneous and heterogeneous cellular compositions. This finding highlights the advantages of three-dimensionality and modularity in approaching the structural and functional properties of the brain compared to classic bidimensional models. Assessing complexity, while the homogeneous model exhibited a notable increase, the heterogeneous one showed a more stable pattern, probably related to the key role of inhibition in regulating activity. In summary, three-dimensionality and modularity are indispensable for accurately replicating the brain’s complexity and richness, while heterogeneity primarily contributes to an expanded repertoire of fragmented network bursts, resulting in nuanced activity patterns and dynamical variability.

In conclusion, we have presented here a three-dimensional model predominantly human-derived, with a minor contribution of rat astrocytes, in which heterogeneity is defined by a controlled mixture of excitatory and inhibitory neurons. We demonstrated its capability to approximate both dynamical richness and the complexity of the brain cortex. Our results open up the possibility to explore the properties and functional mechanisms of the biological system of origin, thus serving as a basis for the development of accurate models of neural diseases in the direction of personalized and precision medicine.

## AUTHOR DECLARATIONS

### Competing interests

The authors declare no competing interests.

### Ethics Approval

We received the Ngn2-positive and Ascl1-positive hiPSCs lines in frozen vials, kindly provided by Prof. Nadif Kasri (Radboud University Medical Centre, the Netherlands). There was initial ethical approval for collection of human cells, and the donors had signed informed consent to participate in the study (Coriell Institute for medical research, GM25256 and KULSTEM iPSC core facility Leuven, Belgium, KSF-16-025). The original source confirmed that the research was conducted in accordance with the principles embodied in the Declaration of Helsinki and in accordance with local statutory requirements. The genetically modified organism (GMO) approval under which the lines have been used is IG22-071. The two lines provided by our collaborators were previously characterized (Mossink, van Rhijn, et al., 2021). The lines were infected, according to a previously published protocol (Frega, Van Gestel, et al. 2017), with lentiviral constructs encoding rtTA combined with Ngn2 (Control line 1) or Ascl1 (Control line 2) to generate doxycycline-inducible excitatory or inhibitory neurons arrays (Mossink, van Rhijn, et al., 2021, Mossink, Verboven, et al., 2021). Both lines were generated from reprogrammed fibroblasts. Control line 1 (C1, healthy 30-years-old female, Ngn2) was reprogrammed via episomal reprogramming (Coriell Institute for medical research, GM25256). Control line 2 (C2, healthy 51-years-old male, Ascl1) was reprogrammed via a non-integrating Sendai virus (KULSTEM iPSC core facility Leuven, Belgium, KSF-16-025). Karyotypes of hiPSCs lines were verified, and hiPSCs lines were tested for pluripotency and genomic integrity based on single nucleotide polymorphism arrays (Mossink, van Rhijn, et al., 2021, Mossink, Verboven, et al., 2021). We declare that the research was conducted in accordance with the principles embodied in the Declaration of Helsinki and in accordance with local statutory requirements. We received rat astrocytes in frozen vials, kindly provided by Dr. Tedesco. The experimental protocol was approved by the European Animal Care Legislation (2010/63/EU), by the Italian Ministry of Health in accordance with the D.L. 116/1992 and by the guidelines of the University of Genova (Prot. 75F11.N.6JI, 08/08/2018).

### Availability of data

The datasets used and analysed during the current study are available from the corresponding author on reasonable request.

### Funding

Work supported by #NEXTGENERATIONEU (NGEU) and funded by the Ministry of University and Research (MUR), National Recovery and Resilience Plan (NRRP), project MNESYS (PE0000006) – A Multiscale integrated approach to the study of the nervous system in health and disease (DN. 1553 11.10.2022).

## Acknowledgments

The authors wish to thank the University of Twente, in particular Dr. Monica Frega, and Dr. Mariateresa Tedesco, for kindly supplying the hiPSCs cultures and rat astrocytes, respectively, and, together with Dr. Roberto Raiteri and Dr. Martina Brofiga, for their kind advice on the experimental procedure and useful discussion. Finally, we thank Dr. Federico Barban and Dr. Marta Carè for kindly supplying *in vivo* recordings.

## Authors’ contributions

**GP**: conception and design, cellular culturing and maintenance, electrophysiological data acquisition, data analysis, interpretation of data, writing — original draft. **GZ**: electrical stimulation data analysis, writing — review and editing. **LC**: morphometric data analysis, writing — review and editing. **CB**: AFM data acquisition and analysis, writing — review and editing. **DDL**: immunofluorescence images acquisition and analysis, writing — review and editing. **MC**: conception, interpretation of data, supervision, validation, review and editing. **SM**: conception, interpretation of data, project administration and resources, supervision, validation. All authors read and approved the final manuscript.

## AI statement

The authors declare that no artificial intelligence–based tools were used at any stage of the study, including design, data analysis, or manuscript preparation.

## Notes

### Competing Interest Statement

The authors have declared no competing interest.

## References

1. Pasca, A. M. et al. Functional cortical neurons and astrocytes from human pluripotent stem cells in 3D culture. Nature Methods 2015 12:7 12, 671–678 (2015).

2. Park, J. et al. A 3D human triculture system modeling neurodegeneration and neuroinflammation in Alzheimer’s disease. Nature Neuroscience 2018 21:7 21, 941–951 (2018).

3. Qian, X., Song, H. & Ming, G. L. Brain organoids: Advances, applications and challenges. Development (Cambridge) 146, (2019).

4. Slanzi, A., Iannoto, G., Rossi, B., Zenaro, E. & Constantin, G. In vitro Models of Neurodegenerative Diseases. Front Cell Dev Biol 8, 534138 (2020).

5. Quadrato, G. et al. Cell diversity and network dynamics in photosensitive human brain organoids. Nature 2017 545:7652 545, 48–53 (2017).

6. Giandomenico, S. L. et al. Cerebral organoids at the air–liquid interface generate diverse nerve tracts with functional output. Nature Neuroscience 2019 22:4 22, 669–679 (2019).

7. Foley, K. E. Organoids: a better in vitro model. Nature Methods 2017 14:6 14, 559–562 (2017).

8. Rabadan, M. A. et al. An in vitro model of neuronal ensembles. 10.1038/s41467-022-31073-1 (2022) doi:10.1038/s41467-022-31073-1.

9. Fitzgerald, K. A., Malhotra, M., Curtin, C. M., O’Brien, F. J. & O’Driscoll, C. M. Life in 3D is never flat: 3D models to optimise drug delivery. Journal of Controlled Release 215, 39–54 (2015).

10. Lancaster, M. A. & Knoblich, J. A. Organogenesisin a dish: Modeling development and disease using organoid technologies. Science (1979) 345, (2014).

11. Birey, F. et al. Assembly of functionally integrated human forebrain spheroids. Nature 2017 545:7652 545, 54–59 (2017).

12. Bousquet, J. & Meunier, J. M. [Organotypic culture, on natural and artificial media, of fragments of the adult rat hypophysis]. C R Seances Soc Biol Fil 156, 65–67 (1962).

13. Pautot, S., Wyart, C. & Isacoff, E. Y. Colloid-guided assembly of oriented 3D neuronal networks. Nature Methods 2008 5:8 5, 735–740 (2008).

14. Tedesco, M. T. et al. Soft chitosan microbeads scaffold for 3D functional neuronal networks. Biomaterials 156, 159–171 (2018).

15. Wang, H. Modeling Neurological Diseases With Human Brain Organoids. Front Synaptic Neurosci 10, 346589 (2018).

16. Costamagna, G., Andreoli, L., Corti, S. & Faravelli, I. iPSCs-Based Neural 3D Systems: A Multidimensional Approach for Disease Modeling and Drug Discovery. Cells 2019*, Vol.* 8, *Page* 1438 **8**, 1438 (2019).

17. Logan, S. et al. Studying Human Neurological Disorders Using Induced Pluripotent Stem Cells: from 2D Monolayer to 3D Organoid and Blood Brain Barrier Models. Compr Physiol 9, 565 (2019).

18. Lee, C. T., Bendriem, R. M., Wu, W. W. & Shen, R. F. 3D brain Organoids derived from pluripotent stem cells: promising experimental models for brain development and neurodegenerative disorders. Journal of Biomedical Science 2017 24:1 24, 1–12 (2017).

19. Yamamoto, H. et al. Impact of modular organization on dynamical richness in cortical networks. Sci Adv 4, (2018).

20. Teller, S. & Soriano, J. Experiments in clustered neuronal networks: A paradigm for complex modular dynamics. AIP Conf Proc 1738, (2016).

21. Yamamoto, H. et al. Modular architecture facilitates noise-driven control of synchrony in neuronal networks. Sci Adv 9, (2023).

22. Parodi, G., Zanini, G., Chiappalone, M. & Martinoia, S. Electrical and chemical modulation of homogeneous and heterogeneous human-iPSCs-derived neuronal networks on high density arrays. Front Mol Neurosci 17, (2024).

23. Mossink, B. et al. Cadherin-13 is a critical regulator of GABAergic modulation in human stem-cell-derived neuronal networks. Molecular Psychiatry 2021 27:1 27, 1–18 (2021).

24. Andolfi, A. et al. A micropatterned thermoplasmonic substrate for neuromodulation of in vitro neuronal networks. Acta Biomater 158, 281–291 (2023).

25. van van Hugte, E. J. H., Schubert, D. & Nadif Kasri, N. Excitatory/inhibitory balance in epilepsies and neurodevelopmental disorders: Depolarizing γ-aminobutyric acid as a common mechanism. Epilepsia 64, 1975–1990 (2023).

26. van Bokhoven, H., Selten, M. & Nadif Kasri, N. Inhibitory control of the excitatory/inhibitory balance in psychiatric disorders. F1000Res 7, (2018).

27. Parodi, G., Brofiga, M., Pastore, V. P., Chiappalone, M. & Martinoia, S. Deepening the role of excitation/inhibition balance in human iPSCs-derived neuronal networks coupled to MEAs during long-term development. J Neural Eng 20, 056011 (2023).

28. Parodi, G. et al. In vitro electrophysiological drug testing on neuronal networks derived from human induced pluripotent stem cells. Stem Cell Research and Therapy 15, 1–14 (2024).

29. Sahara, S., Yanagawa, Y., O’Leary, D. D. M. & Stevens, C. F. The fraction of cortical GABAergic neurons is constant from near the start of cortical neurogenesis to adulthood. Journal of Neuroscience 32, 4755–4761 (2012).

30. Miura, Y. et al. Engineering brain assembloids to interrogate human neural circuits. Nature Protocols 2022 17:1 17, 15–35 (2022).

31. Miura, Y. et al. Generation of human striatal organoids and cortico-striatal assembloids from human pluripotent stem cells. Nature Biotechnology 2020 38:12 38, 1421–1430 (2020).

32. Kim, J. et al. Human assembloid model of the ascending neural sensory pathway. Nature 2025 1–11 (2025) doi:10.1038/s41586-025-08808-3.

33. Colombi, I., Nieus, T., Massimini, M. & Chiappalone, M. Spontaneous and perturbational complexity in cortical cultures. Brain Sci 11, 1453 (2021).

34. Casali, A. G. et al. A theoretically based index of consciousness independent of sensory processing and behavior. Sci Transl Med 5, (2013).

35. D’Andola, M., et al. Bistability, Causality, and Complexity in Cortical Networks: An In Vitro Perturbational Study. Cerebral Cortex 28, 2233–2242 (2018).

36. Zamora-López, G., Chen, Y., Deco, G., Kringelbach, M. L. & Zhou, C. Functional complexity emerging from anatomical constraints in the brain: the significance of network modularity and rich-clubs. Scientific Reports 2016 6:1 6, 1–18 (2016).

37. Zhou, S. & Yu, Y. Synaptic E-I balance underlies efficient neural coding. Front Neurosci 12, 307227 (2018).

38. Denève, S. & Machens, C. K. Efficient codes and balanced networks. Nat Neurosci 19, 375–382 (2016).

39. Costa, E. C., de Melo-Diogo, D., Moreira, A. F., Carvalho, M. P. & Correia, I. J. Spheroids Formation on Non-Adhesive Surfaces by Liquid Overlay Technique: Considerations and Practical Approaches. Biotechnol J 13, (2018).

40. Osaki, T. et al. Complex activity and short-term plasticity of human cerebral organoids reciprocally connected with axons. Nature Communications 2024 15:1 15, 1–13 (2024).

41. Dingle, Y.-T. L. et al. Three-Dimensional Neural Spheroid Culture: An In Vitro Model for Cortical Studies. 10.1089/ten.tec.2015.0135 (2015) doi:10.1089/ten.tec.2015.0135.

42. Spedden, E., White, J. D., Naumova, E. N., Kaplan, D. L. & Staii, C. Elasticity Maps of Living Neurons Measured by Combined Fluorescence and Atomic Force Microscopy. Biophysj 103, 868–877 (2012).

43. Spedden, E. & Staii, C. Neuron Biomechanics Probed by Atomic Force Microscopy. International Journal of Molecular Sciences 2013, Vol. 14, *Pages 16124-16140* **14**, 16124–16140 (2013).

44. Warm, D., Schroer, J. & Sinning, A. Gabaergic Interneurons in Early Brain Development: Conducting and Orchestrated by Cortical Network Activity. Front Mol Neurosci 14, 807969 (2022).

45. Wang, D. D. & Kriegstein, A. R. GABA Regulates Excitatory Synapse Formation in the Neocortex via NMDA Receptor Activation. The Journal of Neuroscience 28, 5547 (2008).

46. Segal, M., Greenberger, V. & Korkotian, E. Formation of dendritic spines in cultured striatal neurons depends on excitatory afferent activity. European Journal of Neuroscience 17, 2573–2585 (2003).

47. Sukenik, N. et al. Neuronal circuits overcome imbalance in excitation and inhibition by adjusting connection numbers. Proc Natl Acad Sci U S A 118, (2021).

48. Connors, B. W. & Gutnick, M. J. Intrinsic firing patterns of diverse neocortical neurons. Trends Neurosci 13, 99–104 (1990).

49. Doorn, N., Voogd, E. J. H. F., Levers, M. R., Putten, M. J. A. M. van & Frega, M. Breaking the burst: Unveiling mechanisms behind fragmented network bursts in patient-derived neurons. Stem Cell Reports 19, 1583–1597 (2024).

50. Bertucci, C., Koppes, R., Dumont, C. & Koppes, A. Neural responses to electrical stimulation in 2D and 3D in vitro environments. Brain Research Bulletin vol. 152 265–284 Preprint at 10.1016/j.brainresbull.2019.07.016 (2019).

51. Tedesco, M. F. M. M. S. P. M. M. P. Interfacing 3D Engineered Neuronal Cultures to Micro-Electrode Arrays: An Innovative In Vitro Experimental Model. Journal of Visualized Experiments 10.3791/53080 (2015) 10.3791/53080.

52. Zanini*, G., Parodi*, G., Chiappalone, M. & Martinoia, S. Investigating the reliability of the evoked response in human iPSCs-derived neuronal networks coupled to Micro-Electrode Arrays. APL Bioeng (2023).

53. Colombi, I., Tinarelli, F., Pasquale, V., Tucci, V. & Chiappalone, M. A simplified in vitro experimental model encompasses the essential features of sleep. Front Neurosci 10, 195002 (2016).

54. Corner, M. A. From Neural Plate to Cortical Arousal—A Neuronal Network Theory of Sleep Derived from in Vitro “Model” Systems for Primordial Patterns of Spontaneous Bioelectric Activity in the Vertebrate Central Nervous System. Brain Sciences 2013*, Vol.* 3, *Pages 800-820* **3**, 800–820 (2013).

55. Corner, M. A., Baker, R. E. & van Pelt, J. Physiological consequences of selective suppression of synaptic transmission in developing cerebral cortical networks in vitro: Differential effects on intrinsically generated bioelectric discharges in a living ‘model’ system for slow-wave sleep activity. Neurosci Biobehav Rev 32, 1569–1600 (2008).

56. Saberi-Moghadam, S., Simi, A., Setareh, H., Mikhail, C. & Tafti, M. In vitro Cortical Network Firing is Homeostatically Regulated: A Model for Sleep Regulation. Scientific Reports 2018 8:1 8, 1–14 (2018).

57. Carè, M. et al. The impact of closed-loop intracortical stimulation on neural activity in brain-injured, anesthetized animals. Bioelectron Med 8, 1–14 (2022).

58. Care’, M. Investigating the impact of novel personalized neurostimulation strategies to promote recovery after brain lesions. (University Of Genoa, 2023).

59. Xiao, Y. et al. The role of inhibition in oscillatory wave dynamics in the cortex. European Journal of Neuroscience 36, 2201–2212 (2012).

60. Mann, E. O. & Paulsen, O. Role of GABAergic inhibition in hippocampal network oscillations. Trends in Neurosciences vol. 30 343–349 Preprint at 10.1016/j.tins.2007.05.003 (2007).

61. Baltz, T., de Lima, A. D. & Voigt, T. Contribution of GABAergic interneurons to the development of spontaneous activity patterns in cultured neocortical networks. Front Cell Neurosci 4, (2010).

62. Severino, F. P. U. et al. The role of dimensionality in neuronal network dynamics. Scientific Reports 2016 6:1 *6*, 1–14 (2016).

63. Frega, M., Tedesco, M., Massobrio, P., Pesce, M. & Martinoia, S. Network dynamics of 3D engineered neuronal cultures: a new experimental model for in-vitro electrophysiology. Scientific Reports 2014 4:1 4, 1–14 (2014).

64. Winter-Hjelm, N. et al. Functional Complexity of Engineered Neural Networks Self-Organized on Structured 3D Interfaces. Small 21, 2410150 (2025).

65. Sharf, T. et al. Functional neuronal circuitry and oscillatory dynamics in human brain organoids. Nature Communications 2022 13:1 13, 1–20 (2022).

66. Degl’Innocenti, E. & Dell’Anno, M. T. Human and mouse cortical astrocytes: a comparative view from development to morphological and functional characterization. Frontiers in Neuroanatomy vol. 17 Preprint at 10.3389/fnana.2023.1130729 (2023).

67. Frega, M. et al. Rapid neuronal differentiation of induced pluripotent stem cells for measuring network activity on micro-electrode arrays. Journal of Visualized Experiments 2017, (2017).

68. Tang, X. et al. Astroglial cells regulate the developmental timeline of human neurons differentiated from induced pluripotent stem cells. Stem Cell Res 11, 743–757 (2013).

69. Shi, Z. et al. Conversion of fibroblasts to parvalbumin neurons by one transcription factor, Ascl1, and the chemical compound forskolin. Journal of Biological Chemistry 291, 13560–13570 (2016).

70. Chiappalone, M., Massobrio, P. & Martinoia, S. Network plasticity in cortical assemblies. Eur J Neurosci 28, 221–237 (2008).

71. Bologna, L. L. et al. Investigating neuronal activity by SPYCODE multi-channel data analyzer. Neural Netw 23, 685–697 (2010).

72. Maccione, A. et al. A novel algorithm for precise identification of spikes in extracellularly recorded neuronal signals. J Neurosci Methods 177, 241–249 (2009).

73. Mossink, B. et al. Human neuronal networks on micro-electrode arrays are a highly robust tool to study disease-specific genotype-phenotype correlations in vitro. Stem Cell Reports 16, 2182–2196 (2021).

74. Carè, M., et al. Towards personalized electroceutical therapy: electrophysiological investigations in a preclinical model of ischemic lesion. Convegno Nazionale di Bioingegneria Padova (2023).

75. Averna, A. et al. Differential Effects of Open- and Closed-Loop Intracortical Microstimulation on Firing Patterns of Neurons in Distant Cortical Areas. Cereb Cortex 30, 2879–2896 (2020).

76. Schindelin, J., et al. Fiji: An open-source platform for biological-image analysis. Nature Methods vol. 9 676–682 Preprint at 10.1038/nmeth.2019 (2012).

77. Cummings, B. S., Wills, L. P. & Schnellmann, R. G. Measurement of Cell Death in Mammalian Cells. Current protocols in pharmacology / editorial board, S.J. Enna (editor-in-chief) … [et al.] 0 12, 10.1002/0471141755.ph1208s25 (2004).

78. H. Hertz. Über die Berührung fester elastischer Körper. Journal für die angewandte Mathematik 92 156–171 (1881).

