## Supplementary Material for "Functional human neurospheroids recapitulate key features of cortical complexity"

### Supplementary Materials

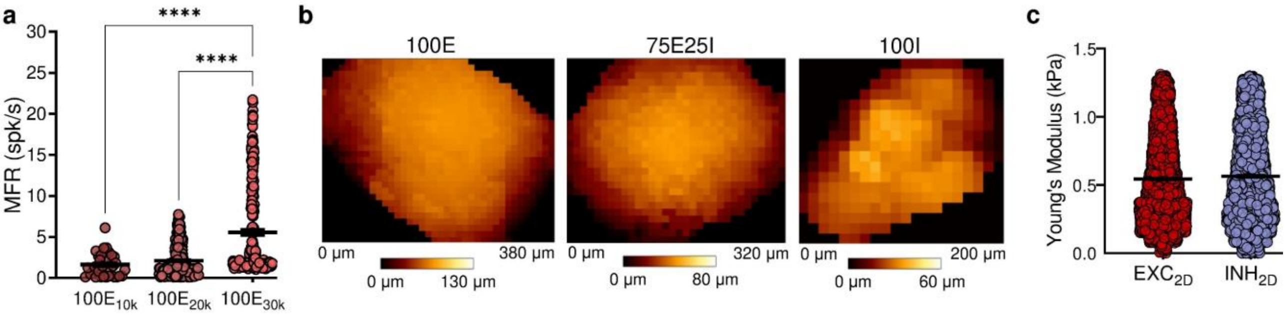


**Supplementary Figure 1: Functional activity, biomechanical topography, and single-cell properties of the model. a)** Mean Firing Rate (MFR) of each channel belonging to $100E_{10k}$, $100E_{20k}$, or $100E_{30k}$ neurospheroids. (Kruskal Wallis' test, * refers to $p < 0.05$, and **** to $p < 0.0001$). **b)** Topography of (from left to right) 100E, 75E25I, and 100I representative neurospheroids. Height values were obtained by force spectroscopy maps at 3 nN applied force. **c)** Young's Modulus of single excitatory (in red) or inhibitory (in blue) cells. In the scatter plots, data are represented with the mean (horizontal line) and the standard error of the mean (whiskers). $N_{EXC2D} = 4183$, $N_{INH2D} = 4383$; Mann-Whitney test with significance level 0.01. Given the large sample size and the minimal effect size observed ($r \approx 0.03$), a more stringent significance threshold of $\alpha = 0.01$ was adopted to reduce the likelihood of detecting trivial differences as statistically significant.

**a**

| Primary Antibody | Host | Dilution | Cat. # | Company |
|---|---|---|---|---|
| NeuN | Guinea Pig | 1:500 | 266 014 | Synaptic System |
| Anti-GABA | Rabbit | 1:200 | A2052 | Sigma-Aldrich |
| MAP2 | Mouse | 1:250 | 13-1500 | Invitrogen |
| GFAP | Rabbit | 1:500 | 173 002 | Synaptic System |

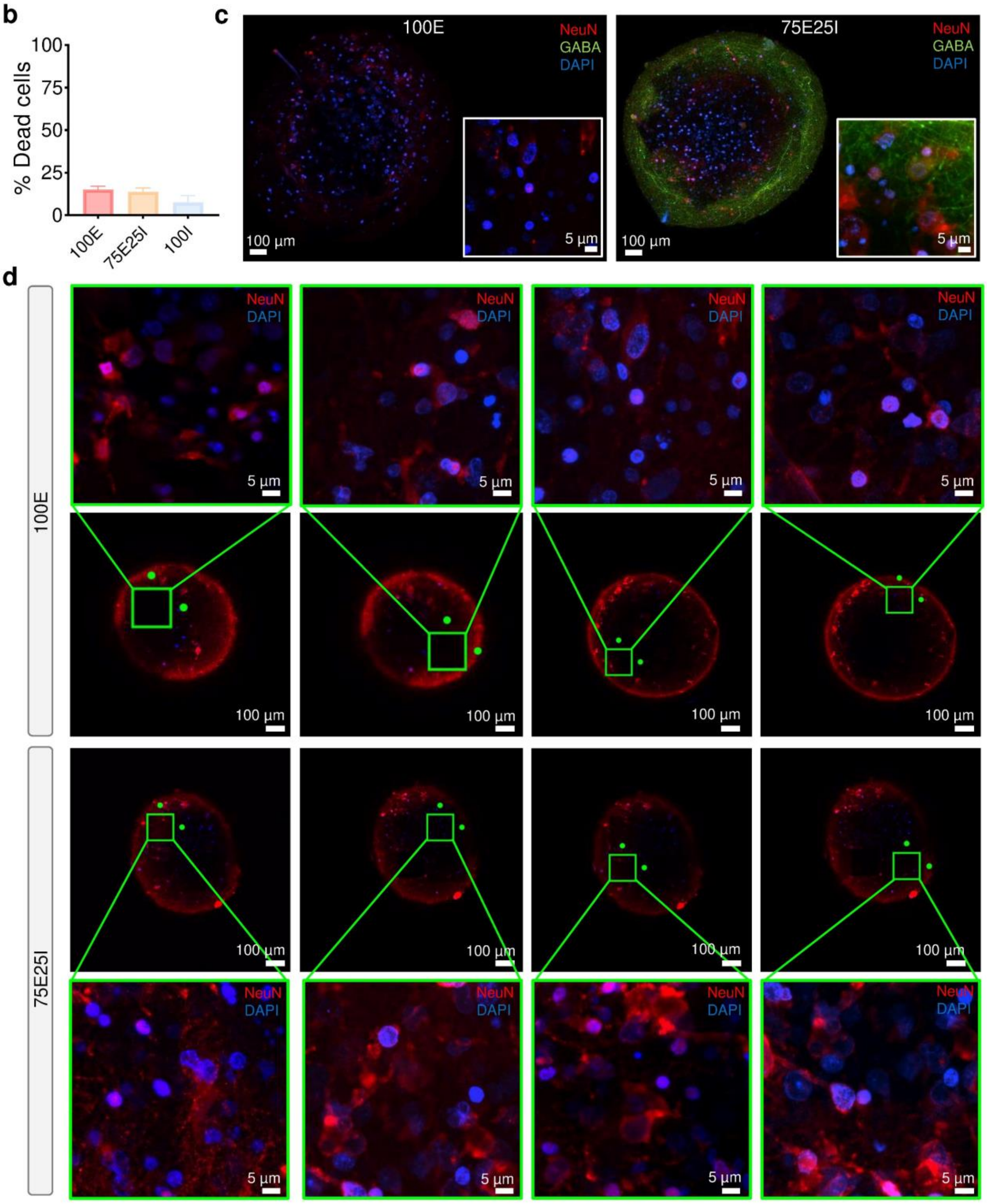


**Supplementary Figure 2: Immunocytochemical characterization and viability assessment of neural spheroids. a)** Table of primary antibodies used for the immunocytochemistry. **b)** Percentage of dead cells at DIV63. **c)** Max projection of z-stacks of 100E (left) and 75E25I (right) spheroids at DIV 70 marked for NeuN (red), GABA (green) and DAPI (blue) and their respective zoom area. **d)** Fluorescence images labelled with NeuN (red) and DAPI (blue) of 100E and 75E25I spheroids and their respective zoom area.

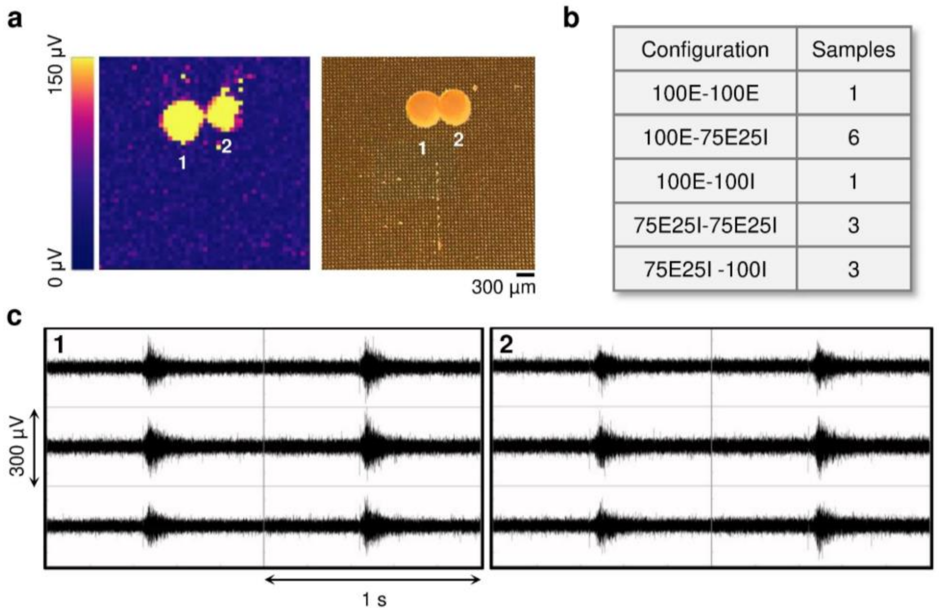


**Supplementary Figure 3: Representative assembloid's activity and dataset. a)** On the left: colour map depicting the amplitude (in µV) of the electrophysiological activity of a representative 3D assembloid in which each pixel represents one channel. On the right: representative image of an assembloid plated on the active area of the high-density MEA. **b)** Dataset of the created assembloids. **c)** Extracellular 1-second signal traces of the representative 3D assembloid in which each box represents one channel.

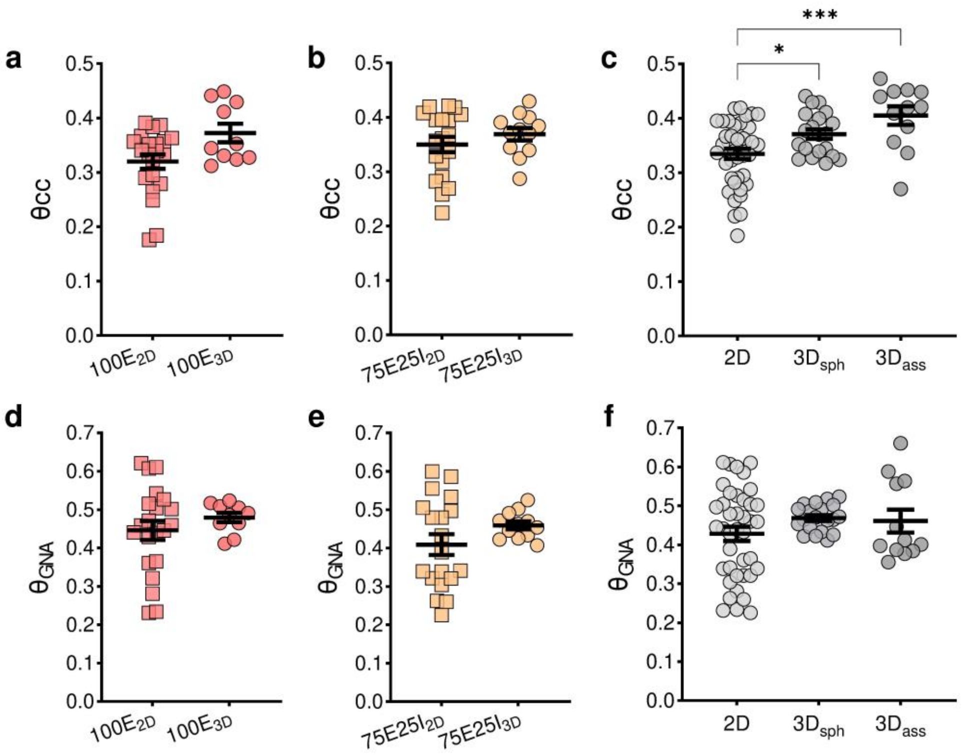


**Supplementary Figure 4: Components of dynamical richness (θ). a-c)** Dynamical richness associated to the variability in the pattern of the spiking activity of 100E configuration (a), 75E25I configuration (b), and related to the model (c). **d-f)** Dynamical richness associated to the variability in the size of the global network activity of 100E configuration (d), 75E25I configuration (e), and related to the model (f). (* refers to $p < 0.05$, and *** to $p < 0.001$; $N_{100E\_2D} = 27$, $N_{100E\_3D} = 12$, $N_{75E25I\_2D} = 23$, $N_{75E25I\_3D} = 14$, $N_{2D} = 50$, $N_{3D\_sph} = 26$, $N_{3D\_ass} = 14$; a), b), d), e) Unpaired t-test; c) and f) One-way Anova test).

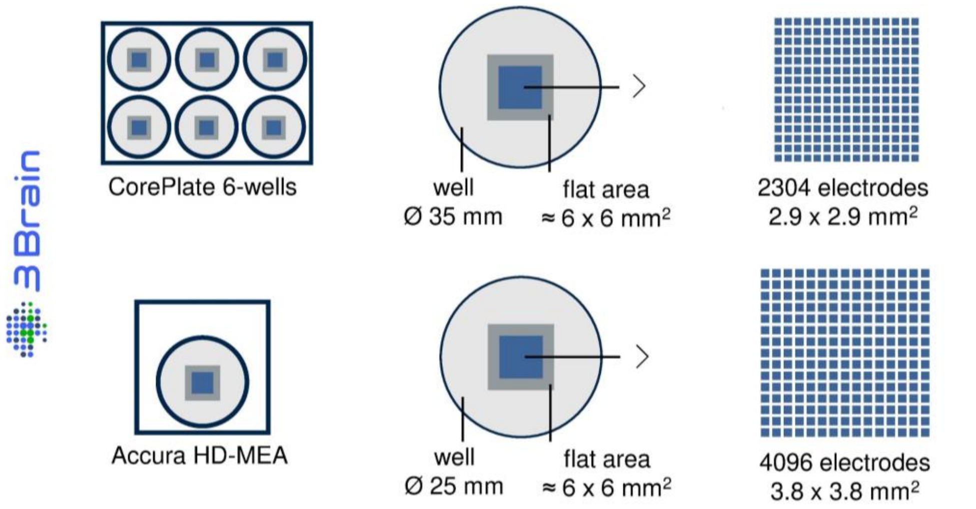


**Supplementary Figure 5: Sketch of the high-density MEAs utilized to record the electrophysiological activity of neurospheroids.** On top: CorePlate devices integrated 2304 recording electrodes characterized by 60 µm in pitch and 25 µm in electrodes' size, arranged in a 2.9 × 2.9 $mm^2$ (48 x 48 electrodes) grid. On bottom: Accura HD devices were characterized by a 3.8 × 3.8 $mm^2$ (64 x 64 electrodes) grid with 21 µm electrodes and 60 µm pitch.

### Captions of Supplementary Movies 1-3

**Supplementary Movie 1:** Rendering 3D of 180 µm of Z-stack of 100E spheroid fixed and immunolabelled for the dendritic marker Map-2 (green), glial marker GFAP (red) and nuclei marker DAPI (Blue).

**Supplementary Movie 2:** Rendering 3D of 123 µm of Z-stack of 75E25I spheroid fixed and immunolabelled for the dendritic marker Map-2 (green), glial marker GFAP (red) and nuclei marker DAPI (Blue).

**Supplementary Movie 3:** Rendering 3D of 110 µm of Z-stack of 100I spheroid fixed and immunolabelled for the dendritic marker Map-2 (green), glial marker GFAP (red) and nuclei marker DAPI (Blue).
